# Development of capsaicin-derived prohibitin ligands to modulate the Aurora kinase A/PHB2 interaction and mitophagy in cancer cells

**DOI:** 10.1101/2024.06.14.598962

**Authors:** Amel Djehal, Claire Caron, Deborah Giordano, Valentina Pizza, Kimberley Farin, Angelo Facchiano, Laurent Désaubry, Giulia Bertolin

## Abstract

Aurora kinase A/AURKA is a serine/threonine kinase overexpressed in a variety of solid and hematological malignancies. In the last decades, clinical trials aiming to counteract the overexpression of AURKA turned out to be largely unsuccessful. Meanwhile, recent discoveries pointed to new functions of AURKA at the subcellular level, including at mitochondria. At this location, AURKA induces organelle clearance by mitophagy acting in complex with the mitophagy mediator LC3, and its inner mitochondrial membrane receptor PHB2. The natural polyphenol xanthohumol was shown to act as a PHB2 ligand, altering the interaction between AURKA and PHB2 and restoring mitochondrial functions in cancer cells. However, its chemical nature prevents its broader use as an anticancer agent.

Using Förster’s Resonance Energy Transfer/Fluorescence Lifetime Imaging Microscopy (FRET/FLIM) in live breast cancer cells, we here explore the effects of alternative PHB ligands in altering the proximity between AURKA and PHB2. Among the already-available compounds, we found that the pungent natural product capsaicin partially alters the AURKA/PHB2 protein-protein proximity. We then synthesized 16 novel capsaicin analogs to enhance the effects of capsaicin. We found that replacing the long hydrophobic acyl moiety with a butyryl one increases the AURKA/PHB2 interaction. Among the capsaicin derivatives carrying this modification, we uncover that compounds **12** and **13** enhance the AURKA/PHB2 proximity. Molecular docking approaches corroborate FRET/FLIM data, and we visualize compounds **12** and **13** in complex with AURKA, PHB2 and LC3. We show that compounds **12** and **13** stabilize the AURKA/PHB2 interaction, and that they can bind to the inhibitory pocket of PHB2 and to the AURKA active site. Finally, we report that compound **13** specifically inhibits AURKA-dependent mitophagy, while leaving the activation of AURKA unaltered at centrosomes.

Together, our data demonstrate that compound **13** is a promising PHB ligand acting on the AURKA/PHB2 interaction. Thanks to its specificity toward the mitochondrial roles of AURKA, it may provide the basis for the development of new anticancer drugs targeting the mitochondrial functions of AURKA.

## Introduction

Mitochondria are in constant equilibrium between turnover and biogenesis. To be able to metabolically sustain an uncontrolled cell proliferation in cancer cells, mitochondria are frequently hijacked to produce significant amounts of ATP^1^. In parallel, damaged mitochondria can be eliminated through a process called mitophagy^2,3^. Mitophagy is controlled by several proteins, including the scaffold protein Prohibitin-2 (PHB2), which binds to the autophagy mediator LC3^4^. PHB2 and its homolog Prohibitin-1 (PHB1) form a heterodimeric complex located on the Inner Mitochondrial Membrane (IMM), and constitute a hub for many signaling pathways beyond mitophagy in cancer cells^5^.

Recently, the overexpression of the cancer-related protein Aurora kinase A/AURKA was shown to induce mitophagy by functionally interacting with PHB2^6^. AURKA regulates a wide variety of mitotic and non-mitotic functions at different subcellular compartments, including mitochondrial dynamics and ATP production^7–11^. We previously showed that the AURKA-dependent phosphorylation of PHB2 on Ser39 induces the formation of an AURKA-LC3-PHB2 tripartite complex^6^. We also showed that this tripartite complex triggers mitophagy in a PARK2/Parkin-independent manner^6^. AURKA-dependent mitophagy is used to eliminate defective mitochondria in cancer cells, while selecting a population of hyperfunctional organelles with high ATP production levels. We also found that the natural polyphenol xanthohumol acts as a PHB2 ligand on the inner membrane of mitochondria to destabilize the AURKA-LC3-PHB2 tripartite complex. By blocking the interaction between AURKA and LC3, treatment with xanthohumol results in mitophagy inhibition. Under these conditions, the AURKA-dependent mitochondrial ATP overproduction is also abolished, thereby linking effective mitophagy and the capacity of mitochondria to increase their ATP production rates^6^. However, the polyphenolic structure of xanthohumol hampers its development as an anticancer agent.

As part of our ongoing programs on the development of new PHB ligands^12–16^, we here screen already-available alternative PHB ligands in their capacity to alter the protein-protein interaction between AURKA and PHB2. We then optimize the activity of the most promising hit – capsaicin - to obtain 16 novel analogs targeting the AURKA/PHB2 interaction. We then illustrate that these ligands differentially affect the capacity of AURKA to interact with PHB2. Using a combination of quantitative fluorescence microscopy and molecular docking, we identify two new PHB ligands dramatically reinforcing the interaction between AURKA and PHB2. By binding both to the AURKA active site and the inhibitory pocket of PHB2, we show that compounds **12** and **13** (N-(4-hydroxy-3-methoxybenzyl)butyramide) behave as inhibitors of this complex at mitochondria. We renamed N-(4-hydroxy-3-methoxybenzyl)butyramide HMBB, and we show that it is specific toward the mitochondrial roles of AURKA. While it blocks AURKA-dependent mitophagy, it does not deactivate the kinase at centrosomes. These features make HMBB a promising target to counteract the effects of AURKA overexpression at mitochondria.

## Results

### Screening of available PHB ligands identifies capsaicin as a compound lowering the interaction between AURKA and PHB2

As an alternative to xanthohumol^6^, we selected three PHB ligands with different chemical structures and with different functional consequences on PHB2 signaling pathways, such as FL3, fluorizoline, and capsaicin^5^ (Figure 1).

**Figure 1.**
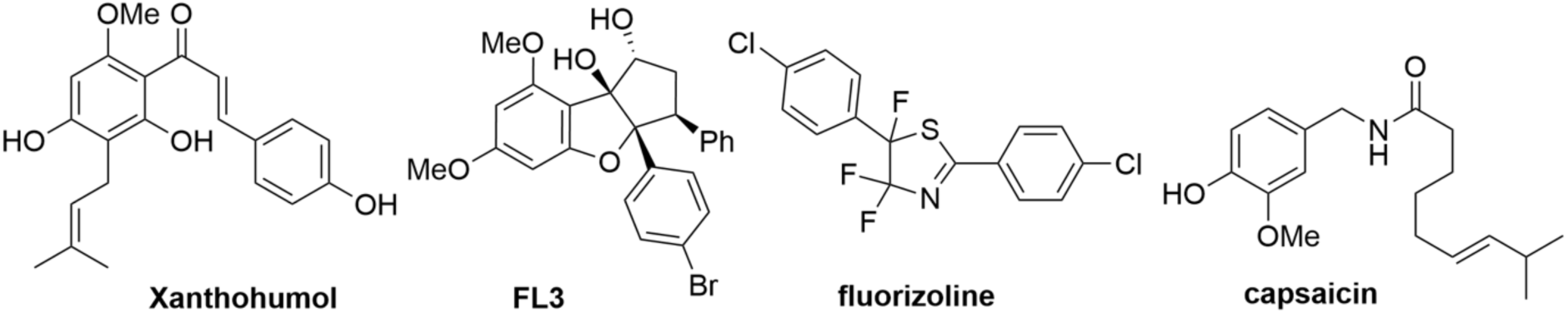
Structure of the PHB ligands xanthohumol, FL3, fluorizoline and capsaicin examined for their effects on the AURKA/PHB2 interaction in breast cancer cells.

We then compared the capacity of FL3, capsaicin and fluorizoline to alter the protein-protein proximity between AURKA and PHB2 using Förster’s Resonance Energy Transfer coupled to Fluorescence Lifetime Imaging Microscopy (FRET/FLIM) in live cells^6,17,18^. In these experiments, AURKA was fused to GFP and was used as a FRET donor, while PHB2 was fused to mCherry and was used as a FRET acceptor. FRET was represented using ΔLifetime, which corresponds to the net difference between the lifetime of AURKA-GFP in the absence (AURKA-GFP + Empty vector, hereby donor-only condition) and in the presence of the acceptor (AURKA-GFP + PHB2-mCherry) (Figure 2). MCF7 breast cancer cells expressing AURKA-GFP in the presence or absence of PHB2-mCherry were incubated for 6h with each of the three compounds, or with the vehicle (DMSO) prior to FRET measurements.

**Figure 2.**
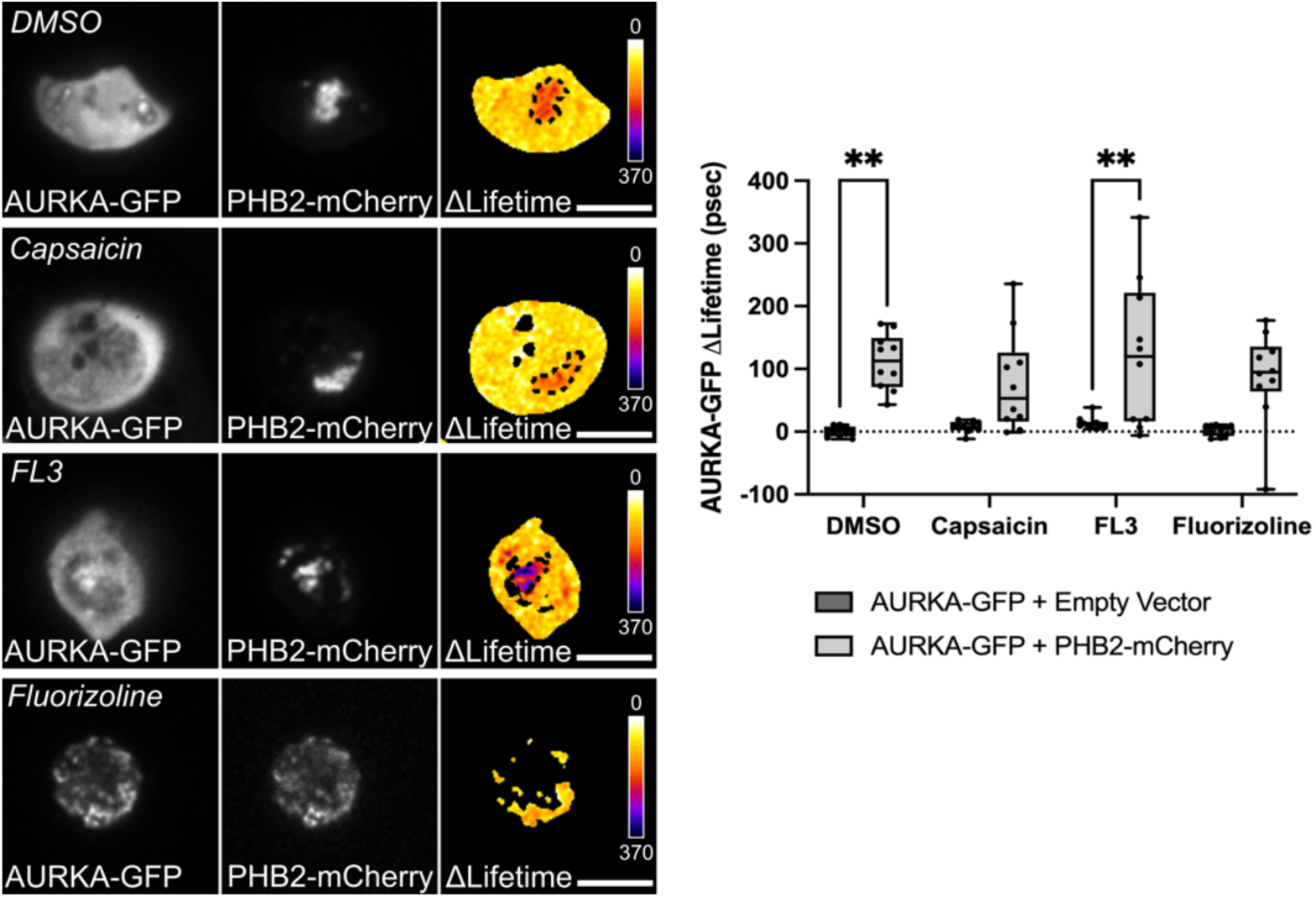
Capsaicin and fluorizoline lower AURKA/PHB2 proximity. FRET/FLIM analyses on MCF7 cells expressing AURKA-GFP with an empty vector (donor-only condition), or with PHB2-mCherry. Cells were treated with vehicle (DMSO), capsaicin, FL3, or fluorizoline (50 µM each). Pseudocolour scale: pixel-by-pixel ΔLifetime. Graph: corresponding ΔLifetime quantifications performed in the dotted area. *n* = 10 cells per condition of one representative experiment (of three). Data extend from the min to the max. Dots indicate individual cells. Scale bar: 10 µm. *\*P*<0.01 against each corresponding “AURKA-GFP + Empty vector” condition. All other comparisons were non-significant.

Treatment with FL3 did not significantly impact the interaction between AURKA and PHB2 in FRET/FLIM analyses. In contrast, treatment with capsaicin or fluorizoline brought the ΔLifetime of AURKA-GFP in the presence of PHB2-mCherry closer to that of their respective donor-only counterparts (Figure 2). This indicates that both compounds significantly lower the interaction between AURKA and PHB2. However, fluorizoline also displayed a significant green autofluorescence that made image-based analyses difficult. Thus, we selected capsaicin as a promising compound for the development of novel PHB ligands that could differentially modulate the AURKA/PHB2 interaction. Capsaicin did not promote a nuclear localization of PHB2 in MCF7 cells, in contrast with a previous report in HeLa cells^19^. Instead, our FRET/FLIM analyses corroborate previous evidence, showing that PHB2 could only be located at mitochondria in MCF7 cells^6^.

Our data show that capsaicin mildly lowers the protein-protein interaction between AURKA and PHB2 at mitochondria. We next synthesized a series of new capsaicin derivatives, and we evaluated their effects on the AURKA/PHB2 interaction.

### Chemical synthesis of novel capsaicin derivatives

After comparing several of methods of acylation, we found that Schotten-Baumann acylation allows the amine of vanillylamine **1a** to be selectively acylated without affecting the phenol as described by Short and coll. (Figure 3A)^20^. With this process, we synthesized 16 new capsaicin derivatives potentially behaving as PHB ligands (Figure 3B). After the synthesis was completed, we then screened these compounds to explore whether they could alter the AURKA/PHB2 interaction at mitochondria.

**Figure 3.**
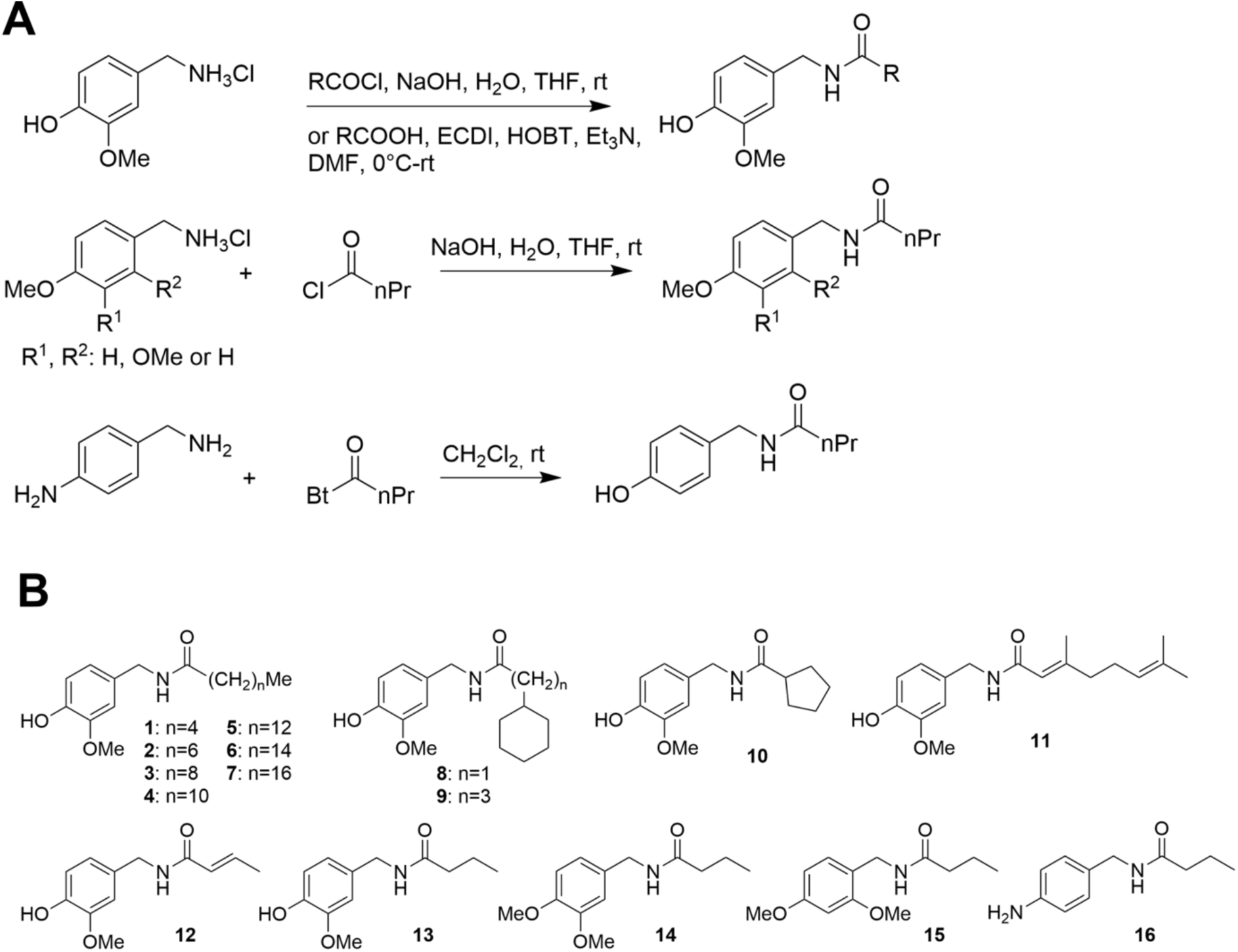
Synthesis reactions (**A**) and structures (**B**) of capsaicin analogs used in the study. Bold numbers below each structure in (B) indicate the number of each compound tested.

### Capsaicin derivatives differentially affect the interaction between AURKA and PHB2

To evaluate the effects of the 16 newly-synthesized capsaicin derivatives (Figure 3B) toward the AURKA/PHB2 interaction, MCF7 cells co-expressing AURKA-GFP and PHB2-mCherry were treated for 6h with each capsaicin derivative or with the DMSO control compound. Their effects on the interaction between AURKA and PHB2 were then evaluated using FRET/FLIM. The dynamic range of a GFP-mCherry chimeric tandem construct shows a maximum of 250 psec of ΔLifetime when compared to a GFP donor-only condition^17^. Given that the AURKA-GFP/PHB2-mCherry pair in the DMSO condition shows a mean ΔLifetime of 120-150 psec *per se*, we explored whether the 16 capsaicin derivatives could induce variations to these mean ΔLifetime values in cells co-expressing AURKA-GFP and PHB2-mCherry.

To this end, we calculated the mean ΔLifetime difference from the AURKA-GFP/PHB2-mCherry pair in the DMSO condition for each novel capsaicin derivative (Table 1). Variations were considered mild when a given compound induced changes in the AURKA-GFP ΔLifetime of +/− 30-50 additional psec compared to what was observed in the AURKA-GFP/PHB2-mCherry pair in the DMSO condition. This corresponds to 12-25% of the maximum ΔLifetime response of a GFP-mCherry tandem^21^. Alternatively, variations were considered significant when a given compound induced changes to the AURKA-GFP ΔLifetime which were superior to +/− 50 additional psec compared to what was observed in the DMSO condition. This is more than 25% of the maximum ΔLifetime response of a GFP-mCherry tandem^21^.

**Table 1.**
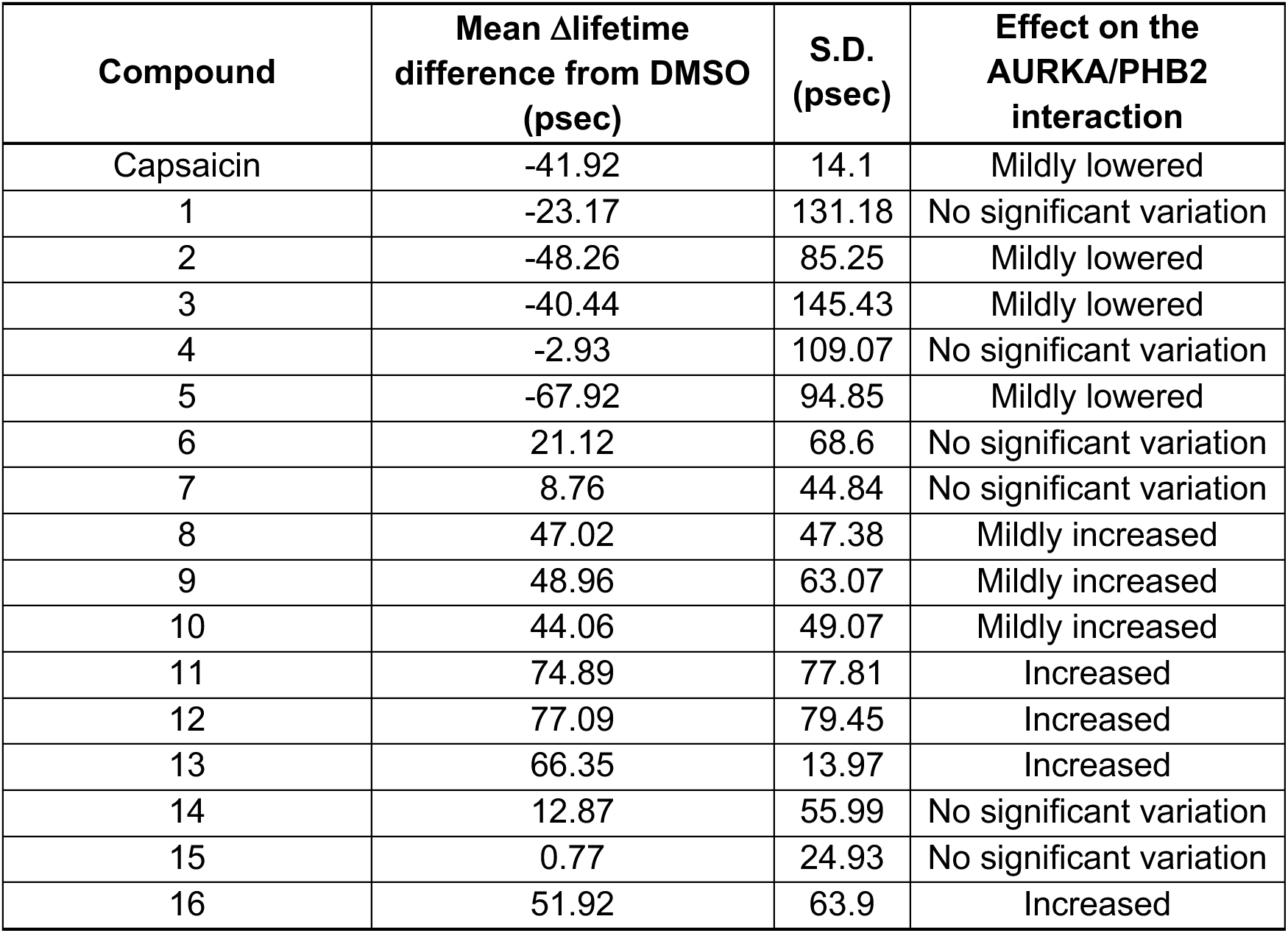
Effect of capsaicin derivatives on the AURKA/PHB2 interaction. Mean ΔLifetime differences and standard deviation (S.D.) of each capsaicin derivative when compared to the vehicle compound, in MCF7 cells co-expressing AURKA-GFP and PHB2-mCherry. Mild variations are between 30 and 50 psec, while variations are considered significant when they are more than 50 psec from the DMSO condition. Capsaicin and all derivatives were used at 50 µM. Results are means from three independent experiments, with *n* = 10 cells per condition in each replicate.

| Compound | Mean ΔLifetime difference from DMSO (psec) | S.D. (psec) | Effect on the AURKA/PHB2 interaction |
| --- | --- | --- | --- |
| Capsaicin | -41.92 | 14.1 | Mildly lowered |
| 1 | -23.17 | 131.18 | No significant variation |
| 2 | -48.26 | 85.25 | Mildly lowered |
| 3 | -40.44 | 145.43 | Mildly lowered |
| 4 | -2.93 | 109.07 | No significant variation |
| 5 | -67.92 | 94.85 | Mildly lowered |
| 6 | 21.12 | 68.6 | No significant variation |
| 7 | 8.76 | 44.84 | No significant variation |
| 8 | 47.02 | 47.38 | Mildly increased |
| 9 | 48.96 | 63.07 | Mildly increased |
| 10 | 44.06 | 49.07 | Mildly increased |
| 11 | 74.89 | 77.81 | Increased |
| 12 | 77.09 | 79.45 | Increased |
| 13 | 66.35 | 13.97 | Increased |
| 14 | 12.87 | 55.99 | No significant variation |
| 15 | 0.77 | 24.93 | No significant variation |
| 16 | 51.92 | 63.9 | Increased |

The replacement of the acyl moiety of capsaicin by various linear and unsaturated fatty acids in compounds **1-7** (6 to 18 carbons) was detrimental to the FRET effect. Indeed, compounds **1**, **4**, **6**, and **7** showed no effect on AURKA-GFP ΔLifetime when AURKA-GFP and PHB2-mCherry were co-expressed. Compounds **2**, **3**, and **5** mildly lowered the AURKA/PHB2 interaction, but with a standard deviation superior to that of capsaicin (Table 1).

Unexpectedly, the introduction of a 5 or 6-membered ring in the hydrophobic acyl chain resulted in an opposite pharmacological activity. Indeed, the constrained analogs **8**, **9**, and **10** mildly increased the interaction between AURKA and PHB2. Of note, these compounds also show a lower degree of experimental variability than the previously-tested compounds in our FRET/FLIM analyses (Table 1). Similar to compounds **8-10**, *N*-vanillylgeranamide/compound **11** displayed some rigidification as well. However, this compound promoted AURKA/PHB2 interaction to a greater extent than the previous analogs (Table 1). Shortening the hydrophobic chain into a crotonic group as in compound **12** also induced significant changes of more than 50 psec in the AURKA-GFP ΔLifetime, thus increasing the interaction between AURKA and PHB2 compared to that observed in the DMSO condition (Table 1). Nevertheless, both compounds **11** and **12** displayed a high standard deviation. Switching to the more flexible butyryl group, as in compound **13,** was another suitable strategy to significantly increase the AURKA/PHB2 interaction. Furthermore, compound **13** displayed a lower standard deviation than compounds **11** or **12** (Table 1). Thus, we chose to modify this analog to explore the vanillyl moiety. The methylation of the phenolic moiety and leading to compounds **14** and **15** left the interaction between AURKA and PHB2 unaltered (Table 1). On the contrary, its replacement by an aniline as in compound 16 increased the interaction between the two proteins (Table 1). This indicates the requirement for a H-bond donor in this position.

To exclude that the capsaicin derivatives could significantly perturb the lifetime of AURKA-GFP *per se,* each compound was evaluated in the donor-only condition where AURKA-GFP was co-expressed with an empty vector (Supplementary Table 1). Previous reports demonstrated that a ΔLifetime difference below 50 psec corresponds to the standard deviation of our FLIM setup^17,18^. Therefore, a difference in the AURKA-GFP ΔLifetime above 50 psec may indicate that the tested ligand intrinsically affects the lifetime of AURKA-GFP, and this beyond the FRET effect observed when AURKA-GFP is co-expressed with PHB2-mCherry. Globally, the majority of the compounds tested showed a non-significant variation of ΔLifetime in the donor-only condition. Nevertheless, we observed that compound **16** shows a slightly higher variability than the other derivatives under these conditions. This suggests that the differences in ΔLifetime observed upon the co-expression of AURKA-GFP and PHB2-mCherry (Table 1) in cells treated with compound **16** could be partially independent of the FRET effect, and could be explained by an intrinsic effect of the compound **16** on the donor lifetime. Therefore, we decided not to use compound 16 for follow-up analyses in living cells and we concentrated

Overall, our FRET/FLIM results show that the newly-generated capsaicin derivatives differentially modulate the interaction between AURKA and PHB2 at mitochondria in live cells. While the majority had either no effect or a mild effect on this interaction, we determined that compounds **11**, **12**, and **13** increase the AURKA/PHB2 protein-protein proximity (Table 1). Among these, compound **13** significantly increases the AURKA/PHB2 interaction with a low standard deviation (Table 1, Fig. 4A-C). This makes this capsaicin derivative the most promising hit among the 16 compounds generated here. Therefore, we further explored its capacity to modulate the AURKA/PHB2 interaction. Washout analyses revealed that compound **13** strongly stabilizes the interaction between the two proteins even after the drug has been removed. Indeed, 3 h washout of the drug still resulted in an increased interaction when compared to control cells co-expressing AURKA-GFP and a mutated, non-fluorescent PHB2-mCherry control construct known as Dark mCherry (mCherry Y72C^22^, Fig. 4D and Supplementary Fig. 1A-C). While AURKA-GFP and PHB2-mCherry show ~ 7% FRET efficiency in control conditions, compound **13** almost doubles this efficiency (~13%) both before and after washout (Supplementary Fig. 1D, E). This further corroborates the role of compound **13** in strengthening this protein/protein interaction. We also observed that compound 13 is equally effective at 50 µM and 10 µM, while its capacity to stabilize the AURKA/PHB2 interaction significantly decreases at 5 µM (Fig. 4E).

**Figure 4.**
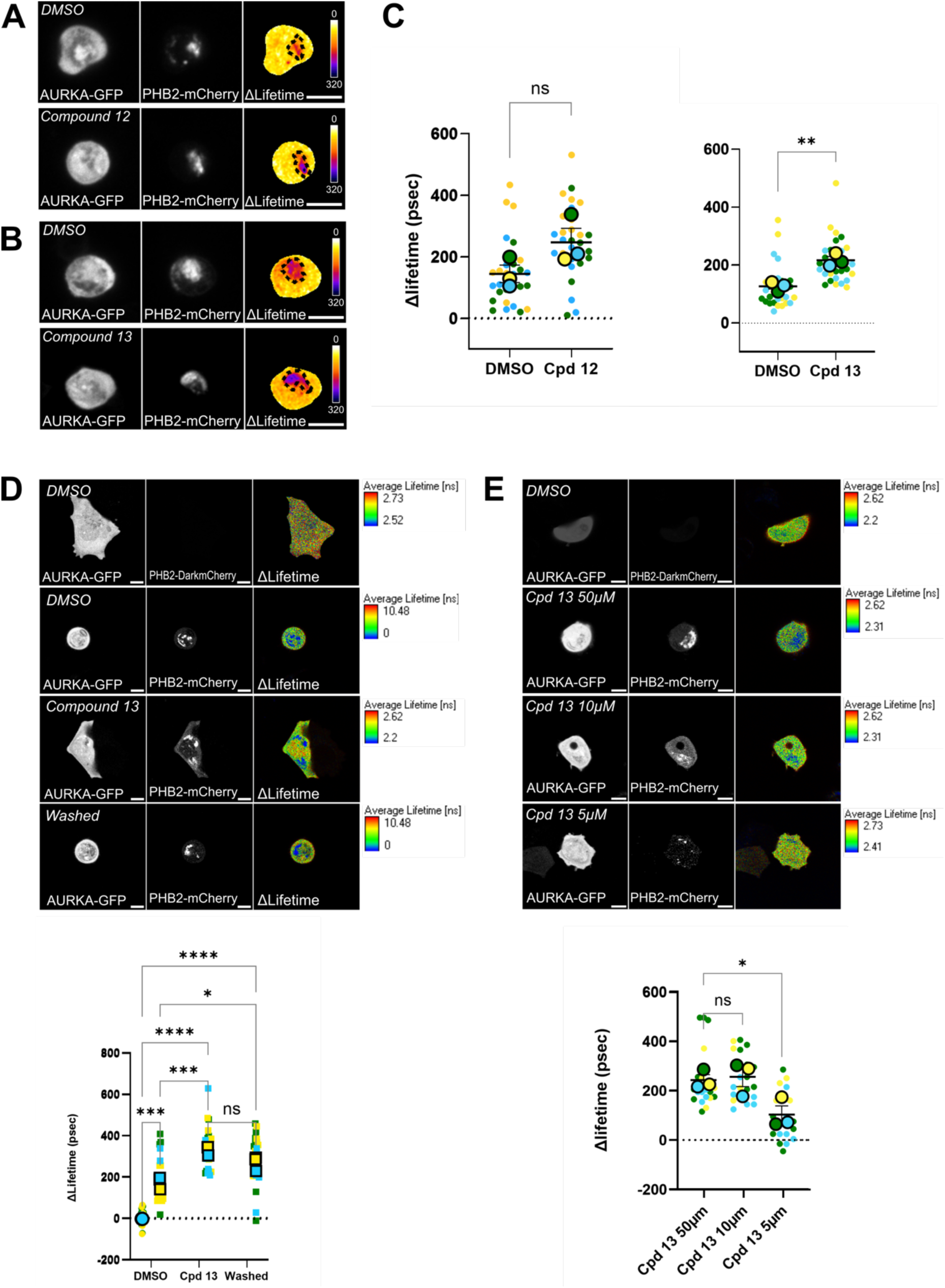
(**A-C**) FRET/FLIM micrographs from MCF7 cells co-expressing AURKA-GFP (donor) and PHB2-mCherry (acceptor). Cells were treated with vehicle (DMSO), compound **12** (**A**, 50 µM) or **13** (**B**, 50 µM). Pseudocolour scale: pixel-by-pixel ΔLifetime. Dotted area: PHB2-rich mitochondrial area. Scale bar: 10 µm. (**C**) ΔLifetime quantifications (in psec) performed in MCF7 cells transfected and treated as in (**A**, **B**). (**D**) Micrographs and TCSPC FRET/FLIM analyses of MCF7 cells co-expressing AURKA-GFP and a non-fluorescent PHB2-Dark mCherry (Donor-only condition, dots) or a fluorescent PHB2-mCherry construct (squares), and treated with DMSO, Compound **13** (50µM, 6h), or Compound **13** (50µM, 6h) followed by 3h washout (“washed” condition). (**E**) Micrographs and TCSPC FRET/FLIM analyses of MCF7 cells co-expressing AURKA-GFP and PHB2-mCherry, and treated with Compound **13** at 50µM, 10 µM, or 5 µm (6h). *n* = 10 cells per condition (small symbols) in each of three experimental replicates. Large symbols indicate mean values per each replicate. Data are means ± S.D. *P*<0.05, *\*\*P*<0.01, *\*\*\*P*<0.001, *\*\*\*\*P*<0.0001. ns: not significant.

Altogether, compound **13** stabilizes the AURKA/PHB2 interaction in MCF7 cells using FRET/FLIM approaches. Therefore, insights at the molecular level are needed to elucidate the binding site of compound **13** to the two proteins.

### Capsaicin and compound 5 induce a molecular clash lowering the AURKA/PHB2 interaction

After observing that compound **13** increases the AURKA/PHB2 interaction, we sought to uncover its binding mechanism to AURKA and PHB2. To this end, we used molecular docking approaches and compared a total of four molecules with opposite FRET/FLIM efficiencies. On one side, we selected capsaicin and compound **5**, which mildly lower the AURKA/PHB2 interaction. On the other side, we selected compounds **12** and **13**, which show the highest FRET/FLIM efficiencies.

First, a reverse docking approach was used to discover possible interesting protein targets for all the selected compounds. Reverse docking results performed by GalaxySagittarius-AlphaFold (Supplementary File 1) highlighted that the four compounds tested were predicted to bind AURKA, even if with different scores. Capsaicin and compound **5** could bind AURKA with a Galaxy best final score of 0.573 and 0.498, respectively. Conversely, compound **12** and **13** could bind AURKA with higher Galaxy best final scores, which were of 0.765 and 0.846, respectively. Interesting, the key capsaicin target Transient Receptor Potential cation channel subfamily V member 1 (TRPV1)^23,24^ could be retrieved only in the presence of capsaicin and compound **5** for the “Binding compatibility prediction” search mode (Table 2). TRPV1 could be retrieved in the list of targets of compound **12**, but only with the “Similarity combined prediction” search mode. TRPV1 also showed a score significantly lower than the score of AURKA, and comparable to that of MAP1LC3 which occupies the lowest position on the list (Table 2). This may suggest a higher specificity of compound **12** and **13**, that could result in a superior affinity and/or selectivity for AURKA.

**Table 2.** Extract of GalaxySagittarius-AF reverse docking results. For each compound, the position (rank) and the score of the proteins of the tripartite complex or of the capsaicin receptor TRPV1 are reported if the target is present among all the hits retrieved.

| Compound | TARGET |  |  |  |  |  |  |  |
| --- | --- | --- | --- | --- | --- | --- | --- | --- |
|  | GALAXY SIMILARITY AlphaFold |  |  |  | GALAXY COMPATIBILITY AlphaFold |  |  |  |
|  | rank | uniprot_id | name | score | rank | uniprot_id | name | score |
| Capsaicin | 20 | O14965 | AURKA | 0,498 | 63 | Q8NER1 | TRPV1 | 0,243 |
|  | 321 | Q8NER1 | TRPV1 | 0,35 | 175 | O14965 | AURKA | 0,222 |
| Compound **5** | 24 | O14965 | AURKA | 0,573 | 457 | Q8NER1 | TRPV1 | 0,238 |
|  | 552 | Q8NER1 | TRPV1 | 0,363 | 780 | O14965 | AURKA | 0,211 |
| Compound<br>**12** | 7 | O14965 | AURKA | 0,765 | 10 | O14965 | AURKA | 0,273 |
|  | 277 | Q8NER1 | TRPV1 | 0,275 |  |  |  |  |
|  | 834 | Q9H492 | MAP1LC3A | 0,211 |  |  |  |  |
| Compound<br>**13** | 4 | O14965 | AURKA | 0,846 | 7 | O14965 | AURKA | 0,273 |

Second, we analyzed the affinity of capsaicin and of compound **5**, **12** and **13** for AURKA and PHB2. To this end, we performed rigid and flexible docking analyses to detect possible relevant binding sites on the activated (pThr288) AURKA structure bound to ADP and Mg^2+^, on the PHB2 model, and on the tripartite AURKA/PHB2/LC3 complex model (Supplementary Table 2). The tripartite AURKA/PHB2/LC3 complex model was built by exploiting rigid protein-protein docking and homology modelling procedures. The best model obtained presents the Ser39 of PHB2^6^ oriented toward the AURKA active site (Supplementary Figure 2). This model also shows the region of LC3 involved in recognition of the LC3-Interacting Region (LIR) binding motif on PHB2 facing the LC3 binding motif YQRL (residues 121-124), and toward the putative LC3 binding motif FNRI (residues 52-55) (Supplementary Figure 2). Binding energy is equal to −12.6 kCal/mol for the AURKA/PHB2 interaction, −10.8 kCal/mol for PHB2/ LC3 interaction, and −1.4 kCal/mol for the AURKA/LC3 interaction. This last value could be due to the very small interface area between AURKA and LC3, which is substantiated by low FRET/FLIM efficiencies in a previous report^6^. Of note, the present model lacks the first 125 residues of AURKA, which are a well-known disordered region. AURKA, LC3 and PHB2 present Z-scores equal to −7.1, −6.6, and −5.91, respectively, in line with the original structures having Z-score values of −7.06, and −6.54 for the crystallographic structure of AURKA and LC3, respectively, and −5.75 for the PHB2 model. The Ramachandran plot of the trimeric complex model obtained presents the following values: 97.8% of dihedral angles in permitted regions of the plot, 4.3% in regions allowed and 0% in generously allowed and disallowed regions.

After generating the model of the tripartite complex (Supplementary Figure 2), we then evaluated the effects of capsaicin and compound **5** on this model. Docking results for compound **5** and capsaicin highlighted a similar behavior of these two compounds (Table 3).

**Table 3.**
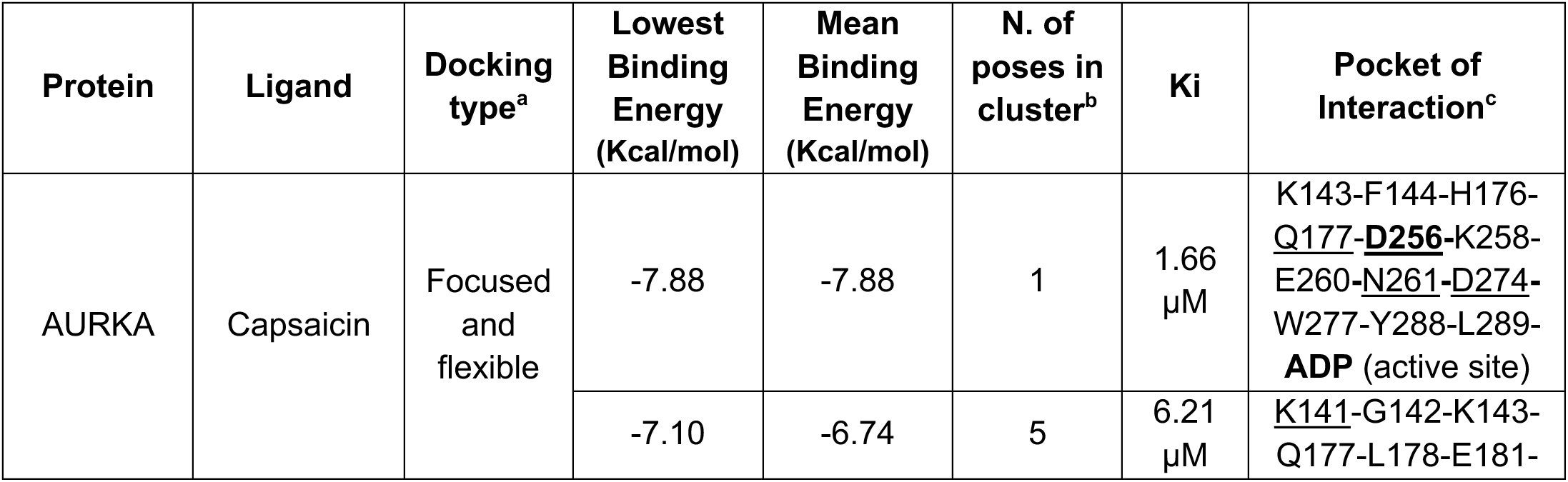
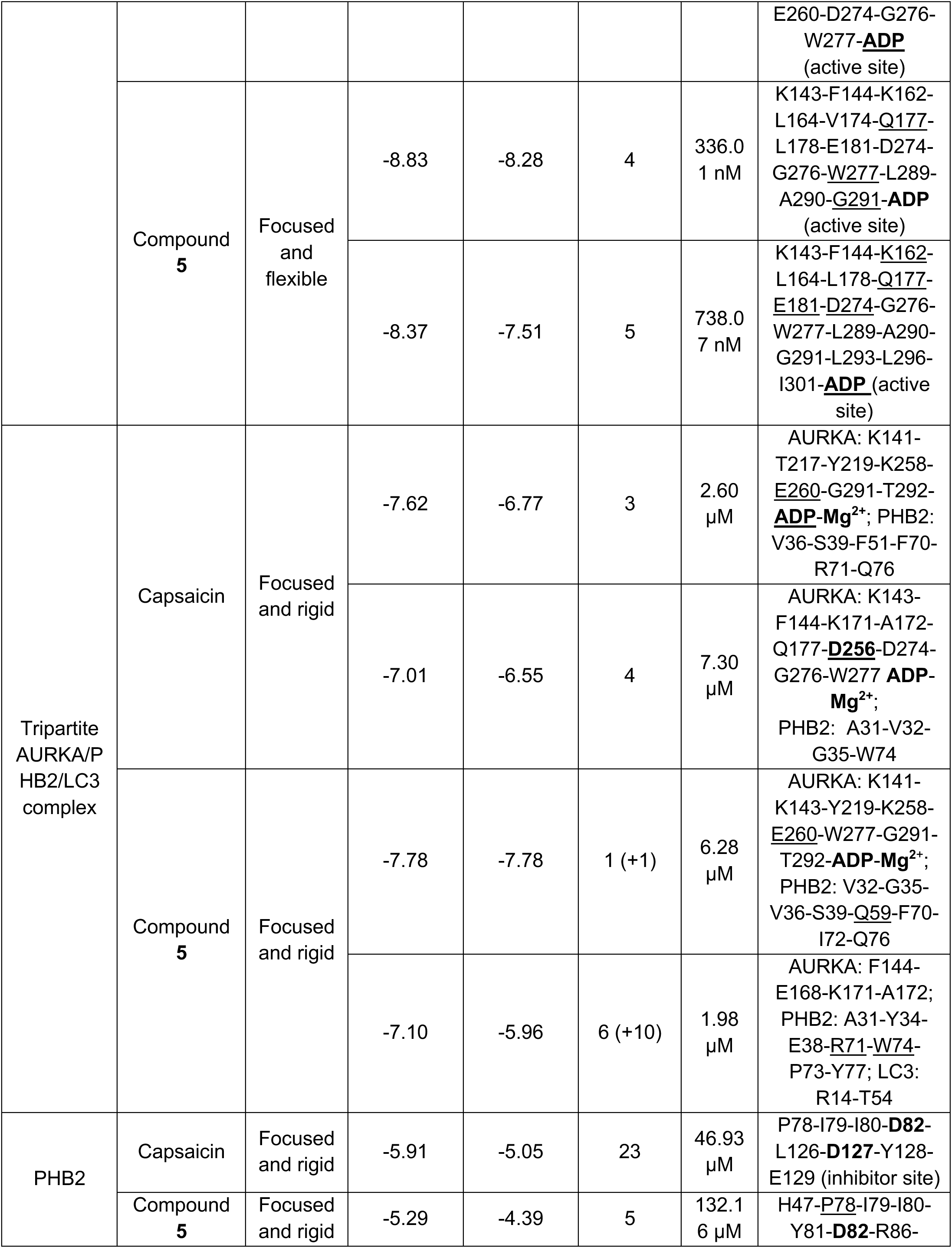
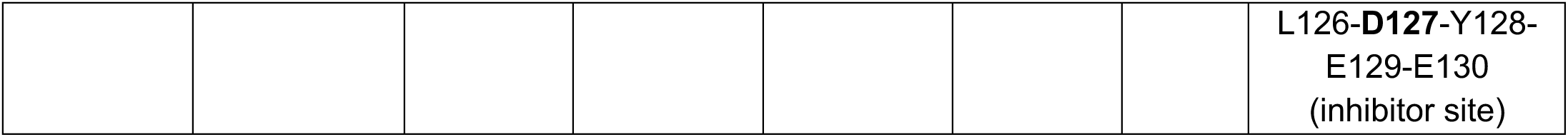
Docking simulation results of capsaicin and compound 5. ^a^: All docking simulations were performed allowing flexibility to the ligand. The annotation “flexible” refers to a docking focused on the AURKA active site, and allowing flexibility not only to the ligand but also to the D256-N261-D274-W277 residues. ^b^: Number of different conformations detected for each ligand in the pocket of interaction, and it is calculated on 100 total poses. ^c^: Residues underlined are involved in H-bond with the ligand, and residues relevant for binding or catalysis are bolded.

| Protein | Ligand | Docking type<sup>a</sup> | Lowest Binding Energy (Kcal/mol) | Mean Binding Energy (Kcal/mol) | N. of poses in cluster<sup>b</sup> | Ki | Pocket of Interaction<sup>c</sup> |
| --- | --- | --- | --- | --- | --- | --- | --- |
| AURKA | Capsaicin | Focused and flexible | -7.88 | -7.88 | 1 | 1.66 μM | K143-F144-H176-Q177-**D256**-K258-E260-N261-D274-W277-Y288-L289-**ADP** (active site) |
| | | | -7.10 | -6.74 | 5 | 6.21 μM | K141-G142-K143-Q177-L178-E181- |
|  |  |  |  |  |  |  | E260-D274-G276-W277-**ADP** (active site) |
|  | Compound **5** | Focused and flexible | -8.83 | -8.28 | 4 | 336.0<br>1 nM | K143-F144-K162-L164-V174-Q177-L178-E181-D274-G276-W277-L289-A290-G291-**ADP** (active site) |
|  |  |  | -8.37 | -7.51 | 5 | 738.0<br>7 nM | K143-F144-K162-L164-L178-Q177-E181-D274-G276-W277-L289-A290-G291-L293-L296-I301-**ADP** (active site) |
| Tripartite AURKA/P HB2/LC3 complex | Capsaicin | Focused and rigid | -7.62 | -6.77 | 3 | 2.60<br>μM | AURKA: K141-T217-Y219-K258-E260-G291-T292-**ADP-Mg<sup>2+</sup>**; PHB2: V36-S39-F51-F70-R71-Q76 |
|  |  |  | -7.01 | -6.55 | 4 | 7.30<br>μM | AURKA: K143-F144-K171-A172-Q177-D256-D274-G276-W277 **ADP-Mg<sup>2+</sup>**; PHB2: A31-V32-G35-W74 |
|  | Compound **5** | Focused and rigid | -7.78 | -7.78 | 1 (+1) | 6.28<br>μM | AURKA: K141-K143-Y219-K258-E260-W277-G291-T292-**ADP-Mg<sup>2+</sup>**; PHB2: V32-G35-V36-S39-Q59-F70-I72-Q76 |
|  |  |  | -7.10 | -5.96 | 6 (+10) | 1.98<br>μM | AURKA: F144-E168-K171-A172; PHB2: A31-Y34-E38-R71-W74-P73-Y77; LC3: R14-T54 |
| PHB2 | Capsaicin | Focused and rigid | -5.91 | -5.05 | 23 | 46.93<br>μM | P78-I79-I80-D82-L126-D127-Y128-E129 (inhibitor site) |
|  | Compound **5** | Focused and rigid | -5.29 | -4.39 | 5 | 132.1<br>6 μM | H47-P78-I79-I80-Y81-D82-R86- |
|  |  |  |  |  |  |  | L126-D127-Y128-E129-E130<br>(inhibitor site) |

Blind and focused, rigid and flexible docking analyses highlighted that compound **5** and capsaicin bind a similar area in the active site of AURKA both with two different possible conformations (Supplementary Table 2). Compound **5** shows interaction with ADP and Mg^2+^ in its best conformation, and binding the AURKA active form. However, it should be noted that only few poses are present in the two clusters, potentially indicating a low selectivity for this area. Both conformations are comparable to the best energetic pose of capsaicin (Supplementary Figure 3). While compound **5** binds exclusively to the cofactors, capsaicin appears to interact with the cofactor and with the active site residue D256. Interestingly, capsaicin binds the AURKA active site with a very low binding energy equal to −7.88 kCal/mol and a constant of inhibition (Ki) of 1.66 µM, and compound **5** does so with an energy value of −8.83 kCal/mol and a Ki of 336.01 nM. All the poses detected suggest a possible inhibitory role of these molecules on the AURKA active site. With this information, we sought to visualize possible steric clashes due to the presence of the ligands. Clashes are conformations occurring when two atoms, which are not covalently bonded to each other, are found to be impossibly close to each other. All the structures of the monomeric proteins bound to the ligands were superimposed on the model of the tripartite complex. The overlap highlighted a possible clash between the ligand – either capsaicin or compound **5** – and PHB2 (Figure 5A), suggesting that the binding of either ligand to AURKA could weaken the AURKA/PHB2 interaction.

**Figure 5.**
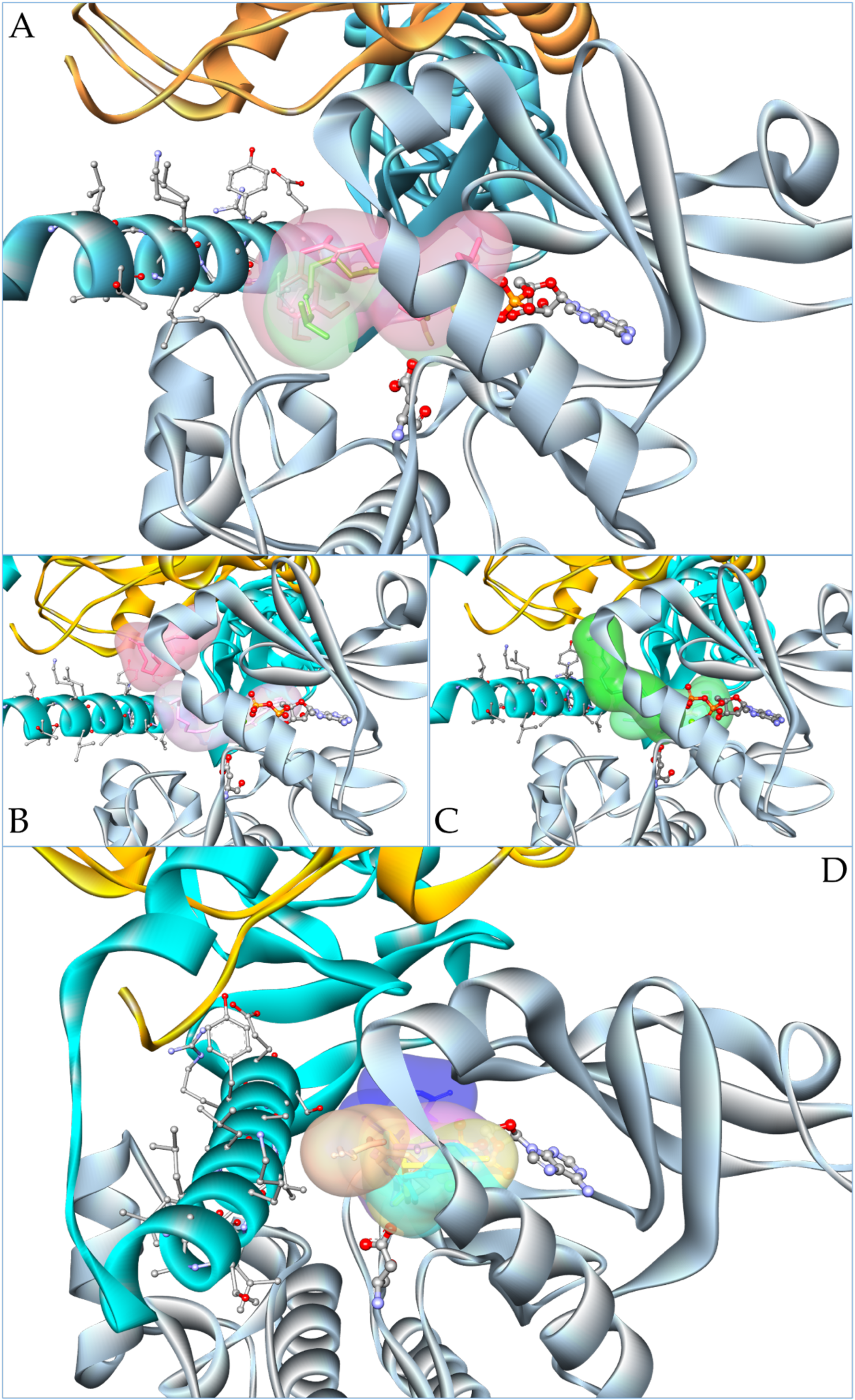
Molecular docking of capsaicin and of compounds **5**, **12** and **13** in complex with the tripartite AURKA/PHB2/LC3 complex model. In all panels, AURKA is represented in grey, PHB2 in cyan and LC3 in orange. Sticks highlight the N-terminal region of PHB2, D256 and ADP in AURKA. (**A**) Binding poses of capsaicin (green sticks and surface) and of compound **5** (pink sticks and surface) to the monomeric active form of AURKA. The overlap shows a clash among the ligands and the N-terminal region of PHB2. (**B**) Results of the docking analysis performed on the tripartite complex with compound **5**. In purple, the best energy binding pose detected is shown, while in pink the representative pose of the most populated cluster is shown. These two poses indicate that compound **5** is forced to move from the pose detected on AURKA alone. (**C**) Results of the docking analysis performed on the tripartite complex with capsaicin. Light green represents the best energy binding pose detected, while the representative pose for the most populated cluster detected is shown in dark green. Capsaicin is forced either to move from the pose detected on AURKA alone or to change its orientation in the pocket. (**D**) Superposition of the conformations of compounds **12** and **13** bound to AURKA alone, with the results of the docking analysis performed on the tripartite complex. Yellow sticks and surface show compound **12** in its AURKA-binding conformation, and in orange the ones detected for the tripartite complex. Cyan sticks and surface show compound **13** in its AURKA-binding conformation. Violet and blue show the binding pose of compound **13** with the best binding energy detected for the tripartite complex and the one representative of the most populated cluster, respectively.

To verify this hypothesis, additional docking simulations were performed on the tripartite complex model. Docking was focused at the interface areas of the three proteins. As expected, compound **5** was found to change its localization on AURKA, with a further 1 kCal/mol loss of its binding energy and reaching values of −7.78 or of −7.10 kCal/mol. Two binding sites were observed to reach the best binding energy with compound 5, and involve the AURKA K141-K143-Y219-K258-E260-W277-G291-T292 residues, the cofactors ADP-Mg2+ and PHB2 V32-G35-V36-S39-Q59-F70-I72-Q76 residues. They also involve AURKA F144-E168-K171-A172, PHB2 A31-Y34-E38-R71-W74-P73-Y77 and LC3 R14-T54 residues (Figure 5B). Similarly to compound 5, capsaicin is able to interact with cofactors in the active site (Figure 5C). However, capsaicin is forced to move into a more dislocated binding site with the best binding energy of −7.62 kCal/mol. Alternatively, capsaicin is forced to adopt an alternative conformation similar to the second one already observed when binding to AURKA, and showing the lowest binding energy equal to −7.10 kCal/mol (Figure 5C).

We then analyzed the effects of capsaicin and compound **5** on PHB2. Docking analyses performed on the PHB2 model were focused on the most organized tertiary structure area of this protein. Results reveal a preference of capsaicin and compound **5** for the inhibitory site of the protein, originally detected when studying the PHB2 inhibitor YL-939^25^. Both capsaicin and compound 5 appear to bind to PHB2 by interacting with the same residues, P78-I79-I80-D82-L126-D127-Y128-E129. The binding energies of these compounds with PHB2 are 5.91KCal/mol and −5.29 kCal/mol for capsaicin and compound **5**. They are higher than the binding energies measured for the two compounds in complex with AURKA, indicating that capsaicin and compound **5** could bind PHB2 with less affinity than for AURKA. However, it should also be noted that the number of poses in the cluster are higher for capsaicin in complex with PHB2 than with AURKA. Despite its lower binding affinity toward PHB2, the number of poses still indicates a good selectivity of capsaicin for the inhibitory site of PHB2.

Overall, capsaicin and compound **5** appear to move from the best energy binding pose to a second pose on AURKA alone. These compounds have a steric hindrance that induces a steric clash at the interface between AURKA and PHB2, and potentially lowering or destabilizing the interaction between these two proteins. The molecular docking data thus substantiate the mild effect of capsaicin and of compound **5** in lowering the AURKA/PHB2 interaction observed with FRET/FLIM experiments.

### Compounds 12 and 13 create a molecular glue between AURKA and PHB2, and they bind to the AURKA active site and the PHB2 inhibitory pocket

Given the previous results on capsaicin and compound **5**, we performed similar binding and docking analyses for compounds **12** and **13**.

Compounds **12** and **13** reach the best binding energy with the active conformation of AURKA, and they appear to specifically bind the substrate-binding site of the kinase (Table 4). Differently from compound **5**, both molecules appear to interact with the D256 residue of the active site and with the cofactors. When observing the structure of the two compounds bound to the tripartite complex, no steric clash between AURKA and PHB2 is observed (Figure 5D). On the contrary, this superposition rather suggests a possible action of compounds 12 and 13 as a bridge between AURKA and PHB2. The docking performed on the complex also revealed that compounds **12** and **13** show a preference for the AURKA binding site (Figure 5D). The binding site is preserved and no clashes occur to alter the binding. The two compounds appear to adapt their orientation in the pocket to preserve the binding and stabilize the complex. Compound **12** preserves its interaction with AURKA Q177, D274, W277 residues, ADP and Mg^2+^. Compound **13** preserves its interaction with AURKA Q177, E181, D256, K258, N261, D274, W277 residues, and with ADP and the two Mg^2+^ ions. Furthermore, the two compounds show an interaction with PHB2. Compound **12** shows interaction with V32, and compound **13** shows interaction with S39, F51, F70, R71, I72, and Q76 in one of the two alternative binding conformations. Binding energies for the compounds binding to the complex or AURKA alone are comparable, with −6.67 kCal/mol and −6.73 kCal/mol for compound **12** bound to the tripartite complex or AURKA alone, and with −6.74 kCal/mol and −6.77 kCal/mol for compound **13** bound to the complex or AURKA alone. This is additional evidence that the binding site does not undergo changes with these two compounds. In addition, the higher number of poses in the cluster suggests that the selectivity of these compounds for the AURKA active is higher than what measured for capsaicin and compound **5**.

**Table 4.** Docking simulation results of compounds 12 and 13. ^a^: All docking simulations were performed allowing flexibility to the ligand. The annotation “rigid” refers to protein residues. ^b^: The number of poses in cluster indicates the number of different conformations detected for each ligand in the pocket of interaction, and it is calculated on a sampling of 100 total poses. ^c^: Residues underlined are involved in the H-bond with the ligand, and residues relevant for binding or catalysis are bolded.

| Protein | Ligand | Docking type<sup>a</sup> | Lowest Binding Energy (kCal/mol) | Mean Binding Energy (kCal/mol) | N. of poses in cluster<sup>b</sup> | Ki | Pocket of Interaction<sup>c</sup> |
| --- | --- | --- | --- | --- | --- | --- | --- |
| AURKA | Compound **12** | Blind and rigid | -6.73 | -6.30 | 6 (+11) | 11.58 μM | F144-L164-L169-Q177- |
|  |  |  |  |  |  |  | L178-E181-<br>**D256**-K258-<br>N261-D274-<br>W277-**ADP**-<br>**Mg<sup>2+</sup>**-<br>**Mg<sup>2+</sup>**(active site) |
|  | Compound<br>**13** | Blind and<br>Rigid | -6.77 | -5.84 | 9 (+9) | 10.86<br>μM | V74-L169-<br>Q177- L178-<br>E181-**D256**-<br>K258-N261-<br>D274-W277-<br>**ADP-Mg<sup>2+</sup>** -**Mg<sup>2+</sup>**<br>(active site) |
| Tripartite<br>AURKA/P<br>HB2/LC3<br>complex | Compound<br>**12** | Focused<br>and<br>Rigid | -6.67 | -5.85 | 3 (+10) | 12.83<br>μM | AURKA: K143-<br>V174-Q177-<br>D274-W277;<br>**ADP-Mg<sup>2+</sup>**;<br>PHB2: V32 |
|  | Compound<br>**13** | Focused<br>and<br>Rigid | -6.74 | -6.45 | 6 | 11.46<br>μM | AURKA: K143-<br>F144-K162-<br>Q177-E181-<br>**D256**-K258-<br>N261-D274-<br>G276-W277-<br>**ADP-Mg<sup>2+</sup>** -**Mg<sup>2+</sup>** |
|  |  |  | -6.51 | -6.19 | 7(+9) | 16.96<br>μM | AURKA: K141-<br>Y219-K258-<br>N261-D274-<br>E260-**ADP**-<br>**Mg<sup>2+</sup>**;<br>PHB2: S39-F51-<br>F70-R71-I72-<br>Q76 |
| PHB2 | Compound<br>**12** | Focused<br>and<br>Rigid | -5.55 | -4.81 | 20 | 85.40<br>μM | I80-Y81-**D82**-<br>R86-L126-<br>**D127**-Y128-E129-E130<br>(inhibitor site) |
|  | Compound<br>**13** | Focused<br>and<br>Rigid | -4.92 | -4.68 | 17 (+5<br>+10) | 248.1<br>4 μM | I80-Y81-**D82**-<br>R86-R88-L126-<br>**D127**-Y128-E129-E130<br>(inhibitor site) |

As performed for capsaicin or compound **5**, we explored the effects of compounds **12** and **13** on PHB2 (Table 4). Both compounds appear to bind the D82-I80-D127 inhibitory site of PHB2, as another PHB2 ligand, YL-939^25^. The two compounds bind to this site with comparable mean binding energy (−4.81 kCal/mol, and −4.68 kCal/mol for compounds **12** and **13**, respectively). The number of poses in each docking cluster is higher for compounds **12** and **13** than for compound **5**. This supports a higher selectivity of the former compounds for the inhibitory pocket than of the latter. The binding energies detectable are lower than those obtained for the AURKA active site. Despite this, the selectivity for the inhibitory pocket of PHB2 is higher, with almost 20 poses out of 100 detected in the same binding area.

Overall, the conformation of compounds **12** and **13** bridges AURKA and PHB2 and brings them close to each other. These compounds potentially act as a molecular glue^26^, stabilizing the interaction between AURKA and PHB2. In addition, docking analyses show that these two compounds not only bind to the AURKA active site but also to the PHB2 inhibitory pocket. Therefore, molecular docking data substantiate the effect of compounds **12** and **13** in strengthening the AURKA/PHB2 interaction observed with FRET/FLIM experiments.

### Compound 13 is specific to the mitochondrial pool of AURKA, and it inhibits AURKA-dependent mitophagy

After modeling the binding of compound **13** to the kinetic pocket of AURKA, we asked whether this molecule was specific to the mitochondrial pool of the kinase. To this end, we first evaluated whether compound **13** could inhibit AURKA activation at alternative known locations of the kinase, such as centrosomes. AURKA activation was evaluated in MCF7 cells expressing the GFP-AURKA-mCherry FRET biosensor^27^. This probe detects the conformational changes of the kinase upon its activation by autophosphorylation on Thr288^28–30^. MLN8237 is widely known to inhibit AURKA activation regardless of its location, including at centrosomes and mitochondria^9,27,31^. As expected, MLN8237 lowered AURKA activation at centrosomes (Fig. 6A). Conversely, compound **13** did not induce any significant FRET perturbation at centrosomes, demonstrating that this molecule is not effective in altering AURKA autophosphorylation at this location (Fig. 6A).

**Fig. 6.**
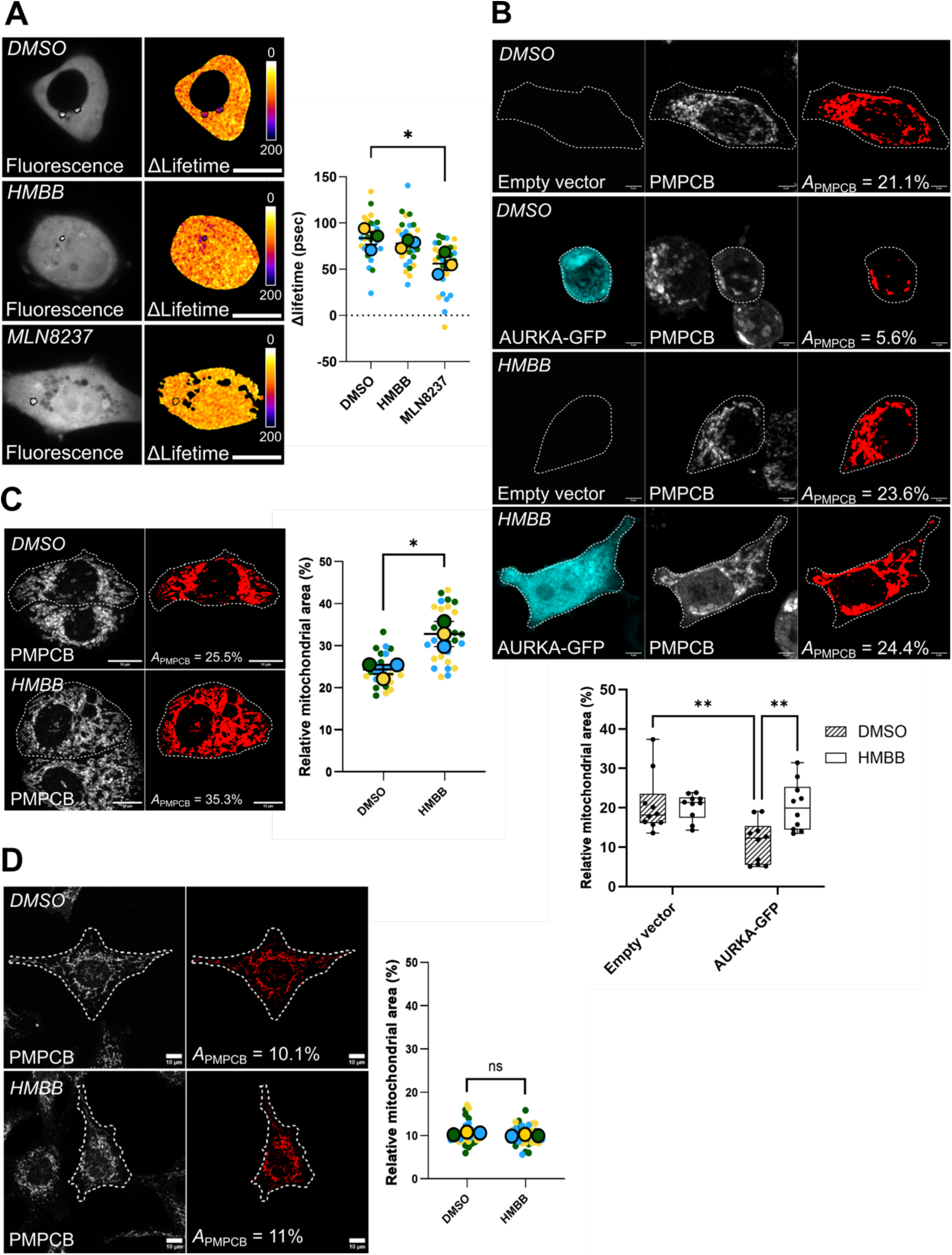
HMBB specifically inhibits AURKA-dependent mitophagy and it does not target the centrosomal pool of the kinase. (**A**) FRET/FLIM images and corresponding quantifications of MCF7 cells expressing the AURKA FRET biosensor and reporting on AURKA activation. Cells were treated with vehicle (DMSO), HMBB (50µM) or the AURKA catalytic inhibitor MLN8237 (100 nM). Pseudocolour scale: pixel-by-pixel ΔLifetime. Dotted area: centrosomes. Scale bar: 10 µm. Graph: ΔLifetime quantifications (in psec) performed in MCF7 cells transfected and treated as indicated. *n* = 10 cells per condition (small dots) in each of three experimental replicates. Large dots indicate mean values for each replicate. Data are means ± S.D. (**B**) Mitochondrial mass shown as the amount of PMPCB staining (threshold mask and corresponding quantification) in MCF7 cells transfected with an empty vector or with AURKA-GFP (pseudocolor cyan), and treated with vehicle (DMSO) or HMBB (50µM) *A*_PMPCB_: mitochondrial area normalized against total cell area (%). *n* = 10 cells per condition from one representative experiment (of three). (**C-D**) Mitochondrial mass shown as the amount of PMPCB staining (threshold mask and corresponding quantification) in T47D cells (**C**) or Hs578T cells (**D**) treated with vehicle (DMSO) or HMBB (50µM). *A*_PMPCB_: mitochondrial area normalized against total cell area (%). *n* = 10 cells per condition from one representative experiment (of three). *n* = 10 cells per condition (small dots) in each of three experimental replicates. Large dots indicate mean values for each replicate. Data are means ± S.D. Scale bar: 5 µm (A-B) or 10 µm (C-D). *\*P*<0.05, *\*\*P*<0.01. ns: not significant.

Compound **13** was renamed as HMBB after its chemical nomenclature N-(4-hydroxy-3-methoxybenzyl)butyramide, and we evaluated its capacity to inhibit AURKA-dependent mitophagy. We previously demonstrated that AURKA-dependent mitophagy is accompanied by a loss of mitochondrial mass, which is calculated by measuring the area covered by the mitochondrial matrix marker PMPCB and normalized over the total cell area^6^. We also showed that this process is PHB2-dependent, and that compounds strengthening the AURKA/PHB2 interaction such as xanthohumol can efficiently block AURKA-dependent mitochondrial clearance^6^.

MCF7 cells expressing AURKA-GFP and treated with DMSO showed a significant mitochondrial mass loss when compared to control cells (Fig. 6B). On the contrary, treatment with HMBB led to an increase in the area covered by PMPCB in cells overexpressing AURKA (Fig. 6B). This is indicative of an increase in mitochondrial mass levels, which are similar to those observed in control cells not overexpressing AURKA. Treatment with HMBB in control cells did not affect mitochondrial mass levels *per se* (Fig. 6B). Therefore, HMBB significantly inhibited AURKA-dependent mitophagy in cells overexpressing the kinase. Accordingly, HMBB increased mitochondrial mass levels in triple-negative T47D breast cancer cells endogenously overexpressing AURKA^9,32^ (Fig. 6C). This is in agreement with the previously-described role of AURKA in regulating mitochondrial mass levels upon overexpression in both MCF7 and T47D cells^6^. Conversely, HMBB had no effect in triple-negative Hs578T cells that have low endogenous AURKA levels (Fig. 6D) ^9,32^. Silencing of *AURKA* or *PHB2* in T47D cells revealed that HMBB cannot rescue mitochondrial mass loss when either component is missing (Supplementary Fig. 4). This further corroborates the specificity of HMBB in AURKA/PHB2-dependent mitophagy.

In conclusion, our results determined that capsaicin is a pharmacological component altering the AURKA/PHB2 protein-protein proximity. Newly-generated capsaicin derivatives showed differential effects on the capacity of AURKA to interact with PHB2, and they either lower or strengthen the interaction in FRET/FLIM experiments. Molecular docking analyses revealed that capsaicin derivatives stabilizing the AURKA/PHB2 proximity can bind to the AURKA/PHB2/LC3 tripartite complex, such as HMBB or compound **12**. These compounds target the AURKA active site and the PHB2 inhibitory pocket, resulting in enhanced binding features compared to molecules lowering the AURKA/PHB2 interaction. Last, we show that HMBB is specific to the mitochondrial pool of AURKA. While it cannot alter its activation at centrosomes, this molecule can efficiently revert AURKA-dependent mitophagy. Overall, we propose that capsaicin derivatives enhancing the AURKA/PHB2 interaction are promising novel inhibitors targeting the mitochondrial roles of AURKA.

## Discussion

Aurora kinase A/AURKA is a hallmark of epithelial and hematological cancers, where it is frequently overexpressed at both the mRNA and protein levels^8^. It is known that AURKA has multiple subcellular locations, including mitochondria^9,10^.

Here, we identified capsaicin as a PHB2 ligand that reduces the AURKA/PHB2 interaction at mitochondria. This compound was previously shown to induce dissociation between PHB and the Adenine Nucleotide Translocator 2 (ANT2, SLC25A5). Proteomics studies suggest that the auto/mitophagy modulator LC3 may interact with ANT2 and in our previous study, we identified ANT2 as an interactor of AURKA using MS/MS-based approaches^9^. In addition, we previously observed that AURKA interacts with LC3 in a PHB2-dependent manner^6^. This raises the interesting possibility that PHB2, AURKA, ANT2 and LC3 may form a quaternary complex in mitochondria.

Capsaicin is known to be cytotoxic in a wide range of cancer cell lines. In addition to PHB2^19^, these effects have been attributed to the action of capsaicin on several targets. These include the neuronal vanilloid receptor TRPV1^23,24^, the Tumor-Associated NADH Oxidase class (tNOX/ENOX2) protein, which is ubiquitously expressed in cancer but not in non-transformed cells^33^, the nuclear lysine-specific demethylase 1A (KDM1A/LSD1)^34^, and the nuclear transcription factor 2 (POU3F2)^35^. Finally, capsaicin was shown to bind to PHB2 and to induce its translocation from the inner mitochondrial membrane to the nucleus in HeLa cells^19^. This also contributes to the induction of apoptosis in human myeloid leukemia cells. Given these multiple targets of capsaicin, it is interesting to note that HMBB and compound **12** are unable to significantly bind to TRPV1^36^. We also observed that capsaicin and compound 5 are the only compounds capable of inducing a cytostatic effect in MCF7 cells, while none of the tested compounds exhibited cytotoxicity (Supplementary Table 3). The lack of overall cytotoxicity exerted by capsaicin and its ligands may suggest low expression levels of canonical capsaicin targets in MCF7 cells. The binding of HMBB and compound **12** is shown when the tripartite AURKA/PHB2/LC3 complex is formed, a condition observable only upon AURKA overexpression^6^. While this makes HMBB and compound **12** two more selective PHB ligands than capsaicin, it also suggests that the detection of cytotoxic effects with these compounds may require AURKA overexpression.

Alternatively, the lack of cytotoxicity may also be explained by the reverse docking results obtained with GalaxySagittarius-AlphaFold predictions. Predictions uncovered that TRPV1 is a preferential target of capsaicin and compound **5**, whereas it was not a preferential target of HMBB or compound **12** (Table 2, Supplementary File 1). This leads to the hypothesis that the cytostatic effect of capsaicin and compound **5** might be related to their activity on TRPV1. With HMBB and compound **12** instead, AURKA stood out as a preferential hit (Table 2, Supplementary File 1). The results on TRPV1 highlight how HMBB and compound **12**, despite being derived from capsaicin, have developed their own selectivity and show a high specificity for other targets such as AURKA. The presence of TRPV1 in the capsaicin list with scores comparable to the best results further corroborates the predictions for the other compounds analyzed. It is important to emphasize that the absence of PHB2 from the GalaxySagittarius list is not related to a potential inability of the compounds to interact with it. Despite the use of GalaxySagittarius in AlphaFold mode, the PHB2 protein model is not present in the reference database. Therefore, evaluating the interaction score between PHB2 and the compounds studied was not possible.

Despite the multiple targets of capsaicin, it is important to note that they are not localized at mitochondria and that the effect of capsaicin at this subcellular compartment was previously unknown. We here provide the first evidence for the capacity of capsaicin and its derivatives to modulate the protein/protein proximity between PHB2 and a cancer-related kinase, such as AURKA, at this compartment. At mitochondria, AURKA was shown to play a variety of roles, including the modulation of mitochondrial morphology, ATP production, and mitochondrial clearance by mitophagy^6,9^. We previously showed that tagging the different proteins with fluorescent tags does not alter their intracellular distributions *per se*^27,9,6,37^. We also showed that the PHB2 ligand xanthohumol globally alters protein-protein interactions within the AURKA-PHB2-LC3 tripartite complex^6^. Indeed, xanthohumol lowers the AURKA/LC3 interaction while concomitantly increasing the AURKA/PHB2 interaction. Altering these tripartite interactions abolishes AURKA-dependent mitophagy and restores normal ATP levels in cancer cells with AURKA overexpression^6^. When looking at the newly-generated capsaicin derivatives, it is interesting to observe that compounds **11**, HMBB, and compound **12** significantly increase the AURKA/PHB2 interaction, as xanthohumol does^6^. Thanks to their low standard deviation compared to that of compound **11** in FRET/FLIM experiments, our data showcase the potential of these capsaicin derivatives for functional studies.

Molecular docking data further corroborate the notion that compounds reinforcing protein-protein interactions by FRET have binding mechanisms compatible with the inhibition of PHB2. All the compounds tested show a preference for a binding site on PHB2 already known to be the target of the YL-939 PHB2 ligand^25^. YL-939 appears to bind PHB2 with two H-bonds involving the D82 and D127 residues, while its distal phenyl group forms hydrophobic interactions with V43 and I80. Mutagenesis experiments performed on D82 and D127 highlighted that these residues are fundamental for the inhibitor binding and reduce the sensitivity to erastin in ferroptosis^25^. Similarly, the binding of capsaicin and compounds **5**, HMBB, and compound **12** to this pocket could exert an inhibitory effect. Moreover, the proximity of the PHB2 inhibition site to the LC3 binding site - YQRL motif, residues 121-124 - may also indicate the involvement of these compounds in modulating the binding of LC3 to PHB2.

Interestingly, capsaicin and compounds **5**, **12** and HMBB all display an interaction with AURKA and show binding energy levels which are better than the ones obtained when interacting with PHB2. However, the different size of the molecules and their different hindrance may explain their differential effects on AURKA. First, they could compete for the binding to Ser39 of PHB2 a residue targeted by AURKA and required for AURKA-dependent mitophagy^6^. As a consequence, the functions of PHB2 within or outside the AURKA/PHB2/LC3 complex could also be compromised. In addition, the modeling of the tripartite complex shows that compound **5** has a steric hindrance for PHB2 accommodation, and this in the two conformations detected. Instead, HMBB and compound **12** accommodate at the interface of AURKA and PHB2. This may give rise to a stabilized AURKA/PHB2 interaction, which is corroborated by FRET/FLIM experiments in living cells.

Docking simulations performed directly on the tripartite complex also underline that HMBB and compound **12** can reach the AURKA active site. These analyses are supported by evidence in living cells, where HMBB can specifically block AURKA-dependent mitophagy. Its inability to target AURKA activation at centrosomes strongly suggests that it is a compound specific to mitochondria, and that it may require the simultaneous presence of AURKA and PHB2 to perform its inhibition. We also show that HMBB inhibits AURKA-dependent mitophagy with the same efficacy as xanthohumol or MLN8237^6^. In this light, the greater bioavailability of capsaicin *in vivo* compared to xanthohumol and the greater mitochondrial specificity of HMBB compared to standard catalytic AURKA inhibitors such as MLN8237 strongly suggest that HMBB and/or capsaicin derivatives with analogous properties may display great selectivity in preclinical or clinical setups.

Our results show that capsaicin derivatives are pertinent chemical tools to alter the formation of the tripartite AURKA/PHB2/LC3 complex both in live cells and at the molecular level. Future studies with a larger number of capsaicin derivatives will elucidate whether the AURKA/PHB2 interaction could be generally considered as a proxy to predict the efficacy of novel pharmacological compounds toward the mitochondrial functions of AURKA and ultimately, the proliferation rates of cancer cells with AURKA overexpression. In conclusion, our study identifies a novel class of bioavailable compounds with the potential to become mitochondrial-specific inhibitors of AURKA.

## Materials and methods

### Cell culture procedures

MCF7 (HTB-22) cells were purchased from the American Type Culture Collection and were kept free from mycoplasma throughout experiments. They were grown in Dulbecco’s Modified Eagle’s Medium (DMEM) already containing 1% L-glutamine (Thermo Fisher Scientific), and supplemented with 10% FBS (Thermo Fisher Scientific) and 1% penicillin-streptomycin (Thermo Fisher Scientific). For all live microscopy experiments, cells were grown at 37°C either in Nunc Lab-Tek Chamber slides (Thermo Fisher Scientific), or in polymer-coated µ-Slide 8 Well high slides (Ibidi). Before proceeding with FLIM imaging, standard growth media was replaced with phenol red-free Leibovitz’s L-15 medium (Thermo Fisher Scientific) supplemented with 20% FBS and 1% penicillin–streptomycin. Plasmid transfections were performed with Lipofectamine 2000 (Thermo Fisher Scientific), according to the manufacturer’s instructions, and incubated for 48h before FLIM analyses. pcDNA3 AURKA-GFP, pcDNA 3.1 empty vector, and pmCherry N1-PHB2 were previously described^6^. PHB2-Dark mCherry (mCherry Y72C^22^) was created with the Quikchange site-directed mutagenesis strategy (Agilent), directly on the pmCherry N1-PHB2 vector using the 5’-CTGTCCCCTCAGTTCATGTGCGGCTCCAAG-3’ and the 5’-CTTGGAGCCGCACATGAACTGAGGGGACAG-3’ forward and reverse primers, respectively. The AURKA conformational biosensor and its donor-only counterpart (GFP-AURKA), originally published in ^27^ were subcloned into a pcDNA 3.1 vector expressing GFP-AURKA-mCherry or GFP-AURKA under the control of the CMV promoter. The siRNA against AURKA was synthesized and purchased from Eurogenetec (sequence: 5’-AUGCCCUGUCUUACUGUCA-3’), as previously described^27^. Allstars negative control (SI03650318) siRNAs and a functionally-validated siRNA against *PHB2* (SI02780918) were purchased from Qiagen. Fluorizoline, FL3, capsaicin, and all capsaicin derivatives were resuspended in DMSO and stored at −20°C for long-term storage. Each compound was used at a final concentration of 50 µM, and incubated for 6h for FRET experiments, or 24h for mitochondrial mass experiments. In dose-response experiments, compound 12 was used at 50 µM, 10 µM, and 5µM, respectively. MLN8237 (Alisertib) was purchased from Euromedex, resuspended in DMSO, and stored at −80°C for long-term storage. It was used at a final concentration of 100 nM and incubated for 6h before imaging.

### FRET/FLIM microscopy

Unless stated otherwise, FLIM analyses were performed as previously described^27^, with a time-gated custom-built system attached to a Leica DMI6000 microscope (Leica) with a CSU-X1 spinning disk module (Yokogawa) and a 63 oil immersion objective (NA 1.4), a picosecond pulsed supercontinuum white laser at 40 MHz frequency (Fianium), and a High-Rate Intensifier (LaVision) coupled to a CoolSNAP HQ2 camera (Roper Scientific). GFP was used as a FRET donor in all experiments, and it was excited at 480 ± 10 nm. Emission was selected using a bandpass filter at 483/35 nm (Semrock). To calculate fluorescence lifetime, five temporal gates with a step of 2 ns each allowed the sequential acquisition of five images covering a total delay time spanning from 0 to 10 ns. The five images were used to calculate the pixel-by-pixel mean fluorescence lifetime according to the equation: *τ* = ∑Δ*τ_i_ ×* I*_i_*/∑I_i_, where Δ*τ_i_* corresponds to the delay time after a laser pulse of the *i*th image acquired. I indicates the pixel-by-pixel fluorescence intensity in each image. The FLIM setup was controlled by the Inscoper Suite solution (Inscoper, France), and GFP lifetime was measured in real-time during acquisition. In all experiments, AURKA-GFP lifetime was calculated by the Inscoper software only when the pixel-by-pixel fluorescence intensity in the first gate was above 2000 gray levels. Raw lifetime values were then extracted by analyzing images post-acquisition with the Fiji software (NIH). To calculate ΔLifetime values, the mean lifetime of the cells in the donor-only condition (AURKA-GFP + Empty vector or AURKA-GFP + PHB2-Dark mCherry for experiments reporting on AURKA/PHB2 interaction; GFP-AURKA for experiments with the GFP-AURKA-mCherry biosensor) was calculated and then used to normalize data in all the analyzed conditions and for each independent experiment as previously reported^38^. FRET efficiency was calculated as previously described^17,27^. FRET/FLIM was also calculated in TCSPC mode (Fig. 4D-E and Supplementary Fig. 1), and GFP lifetime was measured with an inverted SP8 Leica confocal microscope (Manheim, Germany) equipped with a single-molecule detection (SMD) module based on a Picoquant hardware solution (Berlin, Germany), a 470 nm pulsed laser with a 40 MHz repetition rate, a 500/50 nm band pass filter, a single-photon avalanche diode (SPAD) detector, and a 63X oil immersion objective (N.A.= 1.4) as in ^32^. A Picoharp 300 was used for time-correlation and for image reconstruction using scanning signals to recover FLIM images (time-tagged time-resolved (TTTR) method). Lifetime values in each cell were determined by fitting the fluorescence decay with the built-in Symphotime software. This was performed by integrating the signal from the pixels in a region of interest with a single exponential model.

### Immunocytochemistry, confocal microscopy, and mitochondrial mass analyses

Cells were fixed in 4% paraformaldehyde (Euromedex), stained using standard immunocytochemical procedures, and mounted in ProLong Gold Antifade reagent (ThermoFisher Scientific). Mitochondria were stained with a polyclonal rabbit anti-PMPCB (16064–1-AP; Proteintech) used at a 1:500 dilution and a secondary anti-rabbit antibody coupled to Alexa 647 at a 1:5000 dilution (ThermoFisher Scientific). Two-color images of AURKA-GFP and PMPCB-Alexa 647 were acquired with a Leica SP8 inverted confocal microscope (Leica) driven by the Leica Acquisition Suite (LAS) software and a 63x oil-immersion objective (NA 1.4). The excitation/emission wavelengths for GFP were 488 and 525/50 nm and 633 and 650/20 nm for Alexa 647. Mitochondrial mass was determined by calculating the relative mitochondrial area (*A*_PMPCB_) for each image using the PMPCB staining. *A*_PMPCB_ was obtained with the Fiji software (NIH) on maximal projections of confocal images acquired as above. As previously reported^6^, *A*_PMPCB_ was determined as the ratio between the area covered by PMPCB and selected with an automatic threshold mask and the total cell area.

### Cell proliferation and viability

MCF7 cells were cultured in Eagle’s Minimal Essential Medium (EMEM, Sigma-Aldrich) supplemented with 10% FBS (Thermo Fisher Scientific), 2 mM L-Glutamine (Sigma-Aldrich), 1mM sodium pyruvate (Sigma), 100 U/ml penicillin (Sigma-Aldrich) and 100 µg/ml streptomycin (Sigma-Aldrich). Cells were grown at 37°C in NUNC Edge96 Nunclon Delta Surface 96-well plates. All compounds were incubated for 48h prior to cell proliferation and viability experiments, at the following final concentrations: 0.1 µM; 0.5 µM; 1 µM; 5 µM; 10 µM, 50 µM with three independent points measured for each concentration. Only data for 50 µM were reported in Supplementary Table 3. Cellular proliferation was calculated with an IncuCyte S3, driven by the IncuCyte 2023A software, and by measuring cellular confluence. Cellular viability was measured with the WST-1 test (Takara). 450 nm absorbance was recorded using an Envision (Perkin Elmer) plate reader.

### Molecular modelling of capsaicin and capsaicin derivatives, structural models of the AURKA-PHB2-LC3 tripartite complex and molecular docking simulations

The molecular structures of compounds **5**, **12** and **13** were built by Discovery Studio 4.5 (Dassault Systèmes BIOVIA), and minimized by Chimera 1.14^39^. The molecular structure of capsaicin was instead downloaded from PubChem^40^ (CID: 1548943). The putative protein targets of capsaicin and compounds **5-12-13** were estimated with GalaxySagittarius-AF^41^, exploiting both similarity and compatibility mode, and re-ranked using docking. The structures of Aurora kinase A (AURKA) in its active conformation, and of Microtubule-associated protein 1A/1B light chain 3B (LC3) were retrieved from PDB^42^ (PDB code: 5G1X, and 7ELG respectively). For the 3D-structure of Prohibitin-2 (PHB2), the AlphaFold complete model (identifier: AF-Q99623-F1) was used^43^. The H-dock software^44^ was used to create a model of the AURKA-PHB2-LC3 tripartite complex. The complexes obtained were analyzed in terms of compatibility of the interface orientation with the information on the binding site from the literature. Once the most suitable complex was selected, it was used as a template for Modeller9.22^45^ to refine the interactions among the chains of the complex. PDBePISA^46^ was used to calculate the binding energy and the interactions established at the interfaces. ProSA-web^47^ was used to analyze the quality of the model obtained.

Blind and focused docking simulations were performed by AutoDock 4.2.5.1 and AutoDockTools 1.5.6^48^. For blind docking with AURKA active conformation, a 126×126×126 grid box with grid center coordinates equal to (37.908; −15.701; 37.348) was set; spacing was set at 0.397 Å. Focused flexible docking was performed on the AURKA active site to refine the binding energy and the conformation of the compounds in the pocket. A 60×60×60 grid box with center coordinates (39.621; −19.626; 37.357) and a spacing of 0.375 Å was set as default. Four residues - D256-N261-D274-W277 - were set as flexible. This is because D256 is the active site, and the other three residues occupy positions with a relevant role for the access to the active site.

For PHB2, docking was focused on the portion of the protein having a more organized tertiary structure. The greed box was equal to 126×126×126 with center coordinates equal to (−15.757; −2.72; 16.28) and a spacing value of 0.386 Å. Docking simulations performed on the complex with capsaicin and compounds **5**, **12** or **13** were performed focusing the box at the interface of the three molecules with a greed box equal to 126×126×126, a greed center with coordinates (−31.084; −6.4; 22.525), and spacing value of 0.386 Å.

All the docking simulations allowed the flexibility of the ligand. The results obtained were analyzed by AutoDockTools and LigPlot v2.2^49^.

### Statistical analyses

Two-way ANOVA was used to compare the effect of the transfection conditions and of the pharmacological treatments on ΔLifetime (Figure 2, 4D, Supplementary Fig. 1D-E), and on mitochondrial mass loss (Figure 6B, Supplementary Fig. 4). Statistics and summary statistics for biological replicates comparing the effect of the transfection conditions and the pharmacological treatments were performed with *SuperPlotsofData*^50^ (Table 1, Supplementary Table 1). Ordinary one-way ANOVA was used to compare the effect of the pharmacological treatments on the AURKA/PHB2 interaction (Figure 4E) and on the FRET efficiency within the GFP-AURKA-mCherry biosensor (Figure 6A). T-tests were used to compare the effect of transfection conditions on raw and ΔLifetime values (Fig. 4C, Supplementary 1B), and of the pharmacological treatment with compound **13** (HMBB) on mitochondrial mass loss (Fig. 6C-D). Statistics for biological replicates were performed using *SuperPlotsofData* under GraphPad Prism v. 10^51^.

## Author contributions

CRediT taxonomy: Conceptualization (L.D., G.B.) Data Curation (A.D., C.C., D.G., V.P., K.F., A.F., L.D., G.B.), Formal Analysis (A.D., C.C., D.G, V.P., K.F., A.F., L.D., G.B.), Funding Acquisition (L.D., G.B.), Investigation (A.D., C.C., D.G., V.P., K.F., A.F., L.D., G.B.), Methodology (A.F., L.D., G.B.), Project Administration (L.D., G.B.), Resources (L.D., G.B.), Software (C.C., D.G., V.P., A.F., G.B.), Supervision (A.F, L.D., G.B.), Validation (L.D., G.B.), Visualization (C.C., D.G., V.P., A.F., L.D., G.B.), Writing – original, review and editing (C.C., D.G, A.F., L.D., G.B.).

## Acknowledgements

We thank S. Dutertre, X. Pinson and G. Le Marchand at the Microscopy Rennes Imaging Center (MRic, BIOSIT, Biogenouest) for assistance with FLIM experiments. MRic is a member of the national infrastructure France-BioImaging, supported by the French National Research Agency (*ANR-24-INBS-0005 FBI BIOGEN*). We thank the *Plateforme de chimie biologique integrative* (PCBIS, University of Strasbourg) for assistance with cell proliferation and viability experiments. We are grateful to N. Jolivet (IGDR, Rennes) for critical reading of the manuscript, to C. Bertin (IGDR, Rennes) for technical assistance, and to all lab members for helpful comments. This work was supported by the *Centre National de la Recherche Scientifique* (CNRS), the University of Rennes, the *Ligue Contre le Cancer, Comités d’Ille et Vilaine et du Finistère* to G.B. C.C was supported by a PhD fellowship from the *Ligue Nationale Contre le Cancer* (grant no. IP/SC – 17653).

## Data availability

All raw microscopy images are available on Zenodo (DOI: 10.5281/zenodo.17610406).

