## Supplementary Materials and Figures for "Development of capsaicin-derived prohibitin ligands to modulate the Aurora kinase A/PHB2 interaction and mitophagy in cancer cells"

Contains Supplementary Tables 1-3, Supplementary Figures 1-4 and Supplementary Methods. Supplementary File 1 containing the results of GalaxySagittarius-AlphaFold predictions is provided as an additional Excel file.

| Compound | Mean ΔIifetime difference from DMSO (psec) | S.D. (psec) |
|---|---|---|
| Capsaicin | 5.22 | 12.03 |
| **1** | 21.95 | 52.61 |
| **2** | 17.95 | 50.59 |
| **3** | 4.49 | 40.04 |
| **4** | 14.01 | 45.02 |
| **5** | 7.46 | 39.17 |
| **6** | 28.72 | 40.49 |
| **7** | 30.9 | 20.29 |
| **8** | 10.17 | 57.25 |
| **9** | 7.85 | 69.3 |
| **10** | 17.98 | 35.75 |
| **11** | 34.54 | 17.48 |
| **12** | 48.45 | 39.74 |
| **13** | 50.73 | 42.45 |
| **14** | 8.87 | 7.71 |
| **15** | 19.95 | 21.42 |
| **16** | 55.98 | 51.84 |

**Supplementary table 1.** Effect of capsaicin and its derivatives on the AURKA-GFP lifetime. Mean ΔLifetime differences and standard deviation (S.D.) of each capsaicin derivative when compared to the vehicle (DMSO), in MCF7 cells expressing AURKA-GFP and an empty vector. Results are means from three independent experiments, with $n$ = 10 cells per condition in each replicate.

| Flexible docking | | | | | | |
|---|---|---|---|---|---|---|
| **Protein** | **Ligand** | **Lowest Binding Energy (kCal/mol)** | **Mean Binding Energy (kCal/mol)** | **N. of poses in cluster[a]** | **Ki** | **Pocket of Interaction[b]** |
| AURKA (active conformation in complex with $Mg^{2+}$ and ADP) | Capsaicin | -7.88 | -7.88 | 1 | 1.66 μM | K143-F144-H176-Q177-**D256**-K258-E260-N261-D274-W277-Y288-L289-**ADP** (active site) |
| | | -7.10 | -6.74 | 5 | 6.21 μM | K141-G142-K143-Q177-L178-E181-E260-D274-G276-W277-**ADP** (active site) |
| | Compound **5** | -8.83 | -8.28 | 4 | 336.01 nM | K143-F144-K162-L164-V174-Q177-G276-L178-D274-W277-E181-L289-A290-G291-**ADP** |
| | | -8.37 | -7.51 | 5 | 738.07 nM | K143-F144-K162-L164-L178-Q177-E181-D274-G276-W277-L289-A290-G291-L293-L296-I301-**ADP** |
| | Compound **12** | -6.55 | -5.78 | 7 (+10) | 15.84 μM | K143-F144-L169-V174-L178-Q177-**D256**-K258-E260-N261-D274-W277-**ADP** (active site) |
| | Compound **13** | -6.28 | -5.43 | 16 | 24.97 μM | F144-L169-V174-Q177-L178-E181-**D256**-K258-N261-D274-W277-T292-**ADP** (active site) |
| **Rigid Docking** | | | | | | |
| **Protein** | **Ligand** | **Lowest Binding Energy (kCal/mol)** | **Mean Binding Energy (kCal/mol)** | **N. of poses in cluster** | **Ki** | **Pocket of Interaction** |
| AURKA (active conformation in complex with $Mg^{2+}$ and ADP) | Capsaicin | -6.35 | -6.35 | 1(+1) | 21.98 μM | Q177-E181-D274-G276-W277-L289-A290-G291-L296-**ADP-**$\mathbf{Mg^{2+}}$ (active site) |
| | | -5.38 | -4.50 | 5 | 114.69 μM | L289-A290-G291-T292-L293-L296-M300-I301-Y334-Q335-Y338 |
| | Compound **5** | -6.68 | -5.64 | 3 | 12.59 μM | V174-Q177-E181-D274-G276-W277-L289-A290-G291- |

| | | | | | | |
|---|---|---|---|---|---|---|
| | | | | | | L296-M300-I301-**ADP-Mg**$^{2+}$ (active site) |
| | | -6.00 | -5.35 | 3 (+26) | 39.85 µM | L289-A290-G291-T292-L293-D294-L296-M300-I301-E330-A331-N332-T333-Y334-T337 |
| | Compound **12** | -6.73 | -6.30 | 6 (+11) | 11.58 µM | F144-L164-L169-Q177-L178-E181-**D256**-K258- N261-D274-W277-**ADP-Mg**$^{2+}$**-Mg**$^{2+}$(active site) |
| | Compound **13** | -6.77 | -5.84 | 9 (+9) | 10.86 µM | V74-L169- Q177-L178- E181-**D256**-K258-N261-D274-W277-**ADP-Mg**$^{2+}$ **-Mg**$^{2+}$ (active site) |
| PHB2 | Capsaicin | -5.91 | -5.05 | 23 | 46.93 µM | P78-I79-I80-**D82**-L126-**D127**-Y128-E129 (inhibitor site) |
| | Compound **5** | -5.29 | -4.39 | 5 | 132.16 µM | H47-P78-I79-I80-Y81-**D82**-R86-L126-**D127**-Y128-E129-E130 (inhibitor site) |
| | Compound **12** | -5.55 | -4.81 | 20 | 85.40 µM | I80-Y81-**D82**-R86-L126-**D127**-Y128-E129-E130 (inhibitor site) |
| | Compound **13** | -4.92 | -4.68 | 17 (+5 +10) | 248.14 µM | I80-Y81-**D82**-R86-R88-L126-**D127**-Y128-E129-E130 (inhibitor site) |
| Tripartite AURKA/PHB2/LC3 complex | Capsaicin | -7.62 | -6.77 | 3 | 2.60 µM | AURKA: K141-T217-Y219-K258-E260-G291-T292-**ADP-Mg**$^{2+}$; PHB2: V36-S39-F51-F70-R71-Q76 |
| | | -7.01 | -6.55 | 4 | 7.30 µM | AURKA: K143-F144-K171-A172-Q177-**D256**-D274-G276-W277 **ADP-Mg**$^{2+}$; PHB2: A31-V32-G35-W74 |
| | Compound **5** | -7.78 | -7.78 | 1 (+1) | 6.28 µM | AURKA: K141-K143-Y219-K258-E260-W277-G291-T292-**ADP-Mg**$^{2+}$; PHB2: V32-G35-V36-S39-Q59-F70-I72-Q76 |

| | | | | | | |
|---|---|---|---|---|---|---|
| | | -7.10 | -5.96 | 6 (+10) | 1.98 µM | AURKA: F144-E168-K171-A172; PHB2: A31-Y34-E38-R71-W74-P73-Y77; LC3: R14-T54 |
| | Compound **12** | -6.67 | -5.85 | 3 (+10) | 12.83 µM | AURKA: K143-V174-Q177-D274-W277; **ADP-$Mg^{2+}$**; PHB2: V32 |
| | Compound **13** | -6.74 | -6.45 | 6 | 11.46 µM | AURKA: K143-F144-K162-Q177-E181-**D256**-K258-N261-D274-G276-W277-**ADP-$Mg^{2+}$ -$Mg^{2+}$** |
| | | -6.51 | -6.19 | 7(+9) | 16.96 µM | AURKA: K141-Y219-K258-N261-D274-E260-**ADP-$Mg^{2+}$;** PHB2: S39-F51-F70-R71-I72-Q76 |

**Supplementary Table 2**. Flexible and rigid docking simulations of capsaicin and compounds **5**, **12** and **13**.

[a]: The number of poses in cluster indicates the number of different conformations detected for each ligand in the pocket of interaction, and it is calculated on a sampling of 100 total poses.

[b]: Residues underlined are involved in H-bond with the ligand, and residues relevant for binding or catalysis are bolded.

| Compound | Confluence (%) | Viability (%) |
|---|---|---|
| Capsaicin | 73 ± 3% | 107 ± 4% |
| **5** | 75 ± 2% | 94 ± 4% |
| **12** | 99 ± 6% | 105 ± 2% |
| **13** | 95 ± 4% | 110 ± 4% |

**Supplementary Table 3.** Effect of capsaicin and compounds **5**, **12** and **13** on cell confluence and viability, after 48h of treatment at 50 μM in MCF7 cells. Values are means ± S.D.

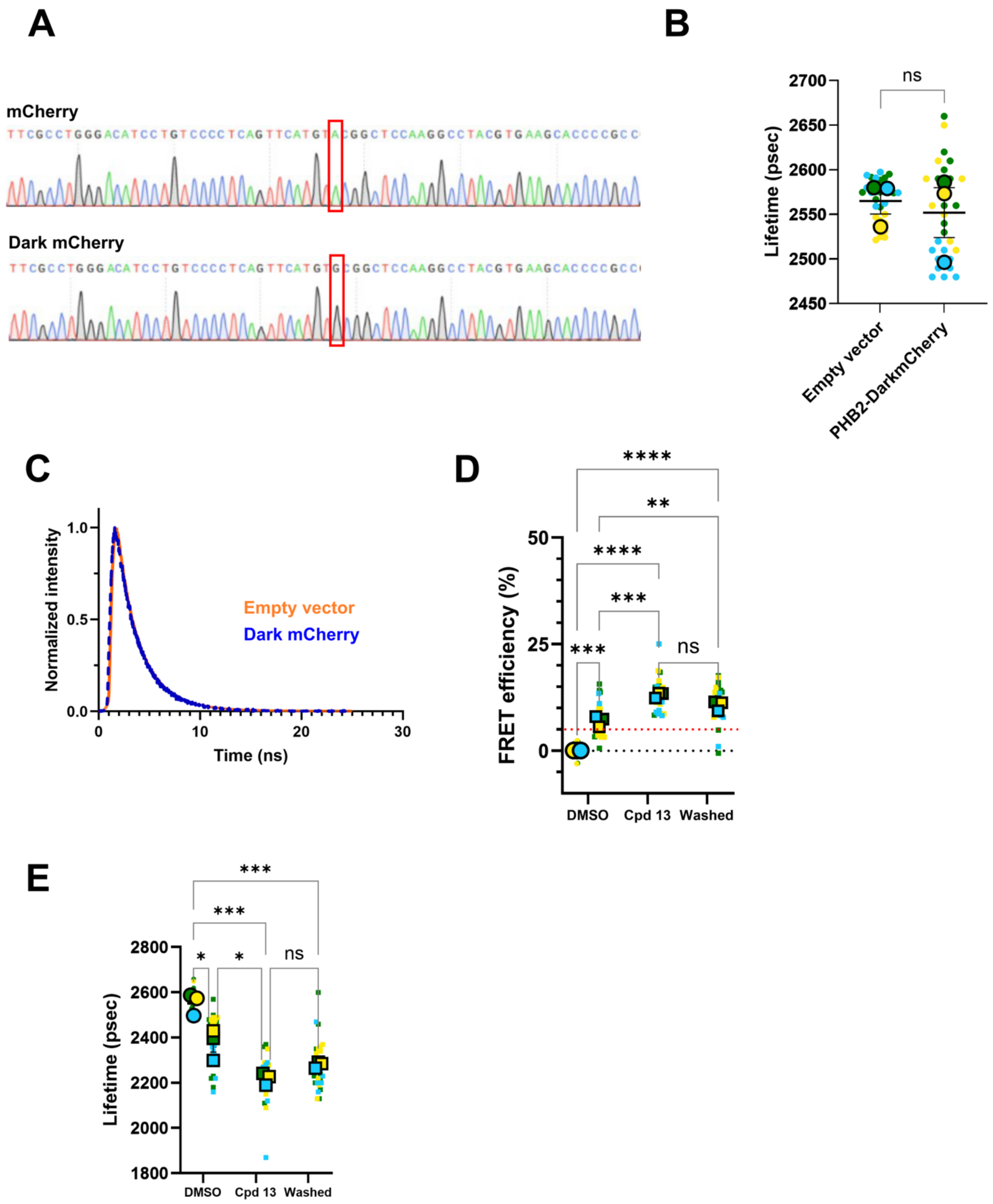


**Supplementary Figure 1**: (**A**) Chromatograms illustrating the Y72C point mutation (red box) on the mCherry sequence to create a Dark mCherry variant. (**B**) AURKA-GFP raw lifetime values (psec) in MCF7 cells co-expressing an empty vector or a non-fluorescent PHB2-Dark mCherry construct. *n* = 10 cells per condition (small dots) in each of three experimental replicates. Large dots indicate mean values for each replicate. (**C**) Representative fluorescence lifetime decay profiles measured by TCSPC in MCF7 cells co-expressing AURKA-GFP with an empty vector (orange profile) or with PHB2-Dark mCherry (blue profile). (**D-E**) FRET efficiency (%, **D**) and corresponding raw lifetime values (**E**) obtained from MCF7 cells co-expressing AURKA-GFP and a non-fluorescent PHB2-Dark mCherry (Donor-only condition, dots) or a fluorescent PHB2-mCherry construct (squares), and treated with DMSO, Compound **13** (50µM, 6h), or Compound **13** (50µM, 6h) followed by 3h washout (“washed”). Red dashed line: 5% net difference between the Donor-only condition (AURKA-GFP + PHB2-Dark mCherry) and the AURKA-GFP + PHB2-mCherry condition in cells treated with DMSO. Data are means ± S.D. $P<0.05$, **$P<0.01$, ***$P<0.001$, ****$P<0.0001$. ns: not significant.

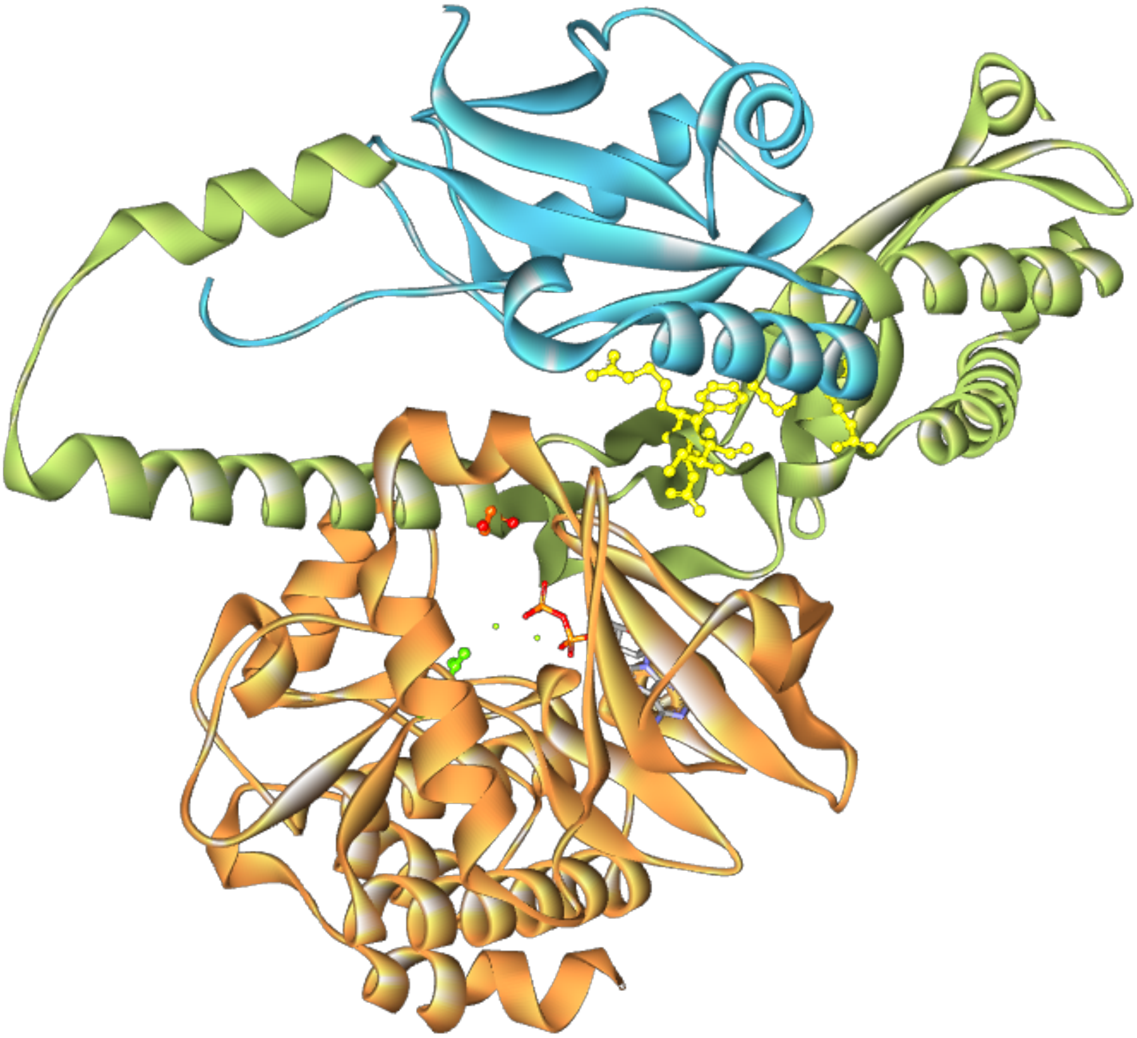

**Supplementary Figure 2**: Best molecular docking model obtained of the tripartite AURKA/PHB2/LC3 complex. PHB2 is represented in green solid ribbon, AURKA in orange, while LC3 is shown in cyan. The model displays Ser39 of PHB2[6] (orange balls and sticks) oriented towards the AURKA active site (ADP is shown in sticks, while the catalytic D245 residue is highlighted in green balls and sticks). LC3 faces the LIR binding motif YQRL (residues 121-124) on PHB2, and a putative LIR binding motif FNRI (residues 52-55) on PHB2. Both LIR motifs are highlighted in yellow balls and sticks.

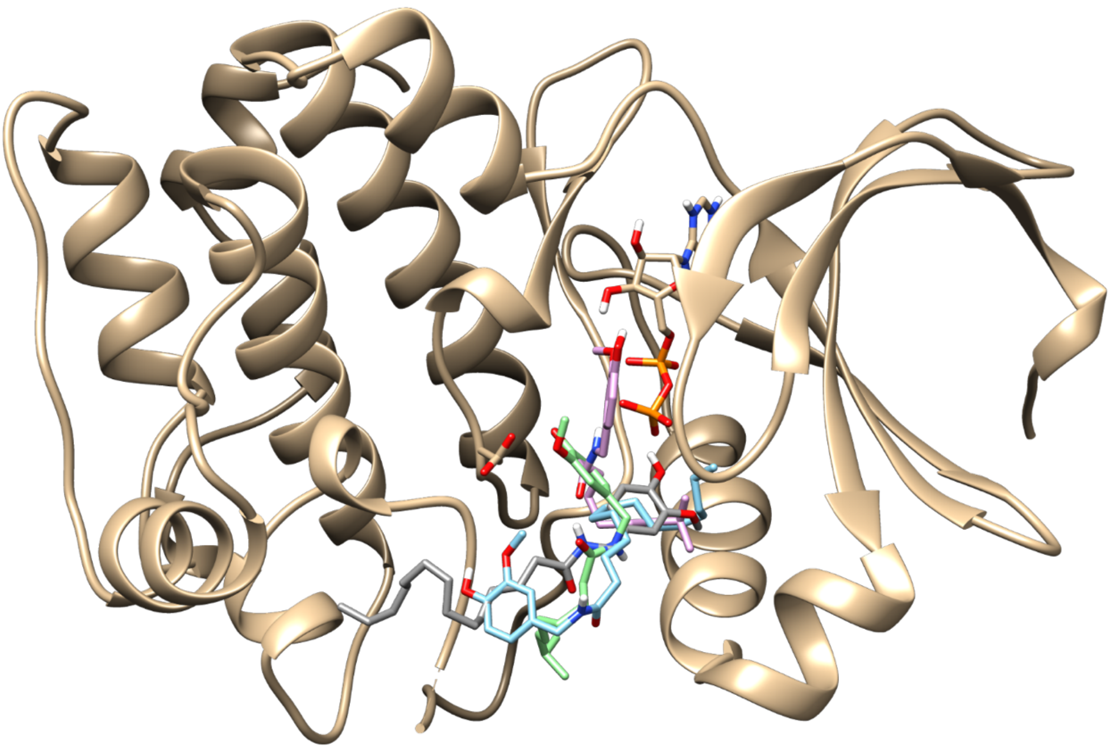

**Supplementary Figure 3**: Comparison of the binding poses adopted by compound **5** and capsaicin in the AURKA active site. Beige ribbons: AURKA; beige sticks: ADP and D256. Green and pink sticks: first and second best energetic conformation of capsaicin; Cyan and grey: first and second best energetic conformation adopted by compound 5. Capsaicin reaches a conformation that allows a better interaction with the catalytic site, in particular with D256.

**A**

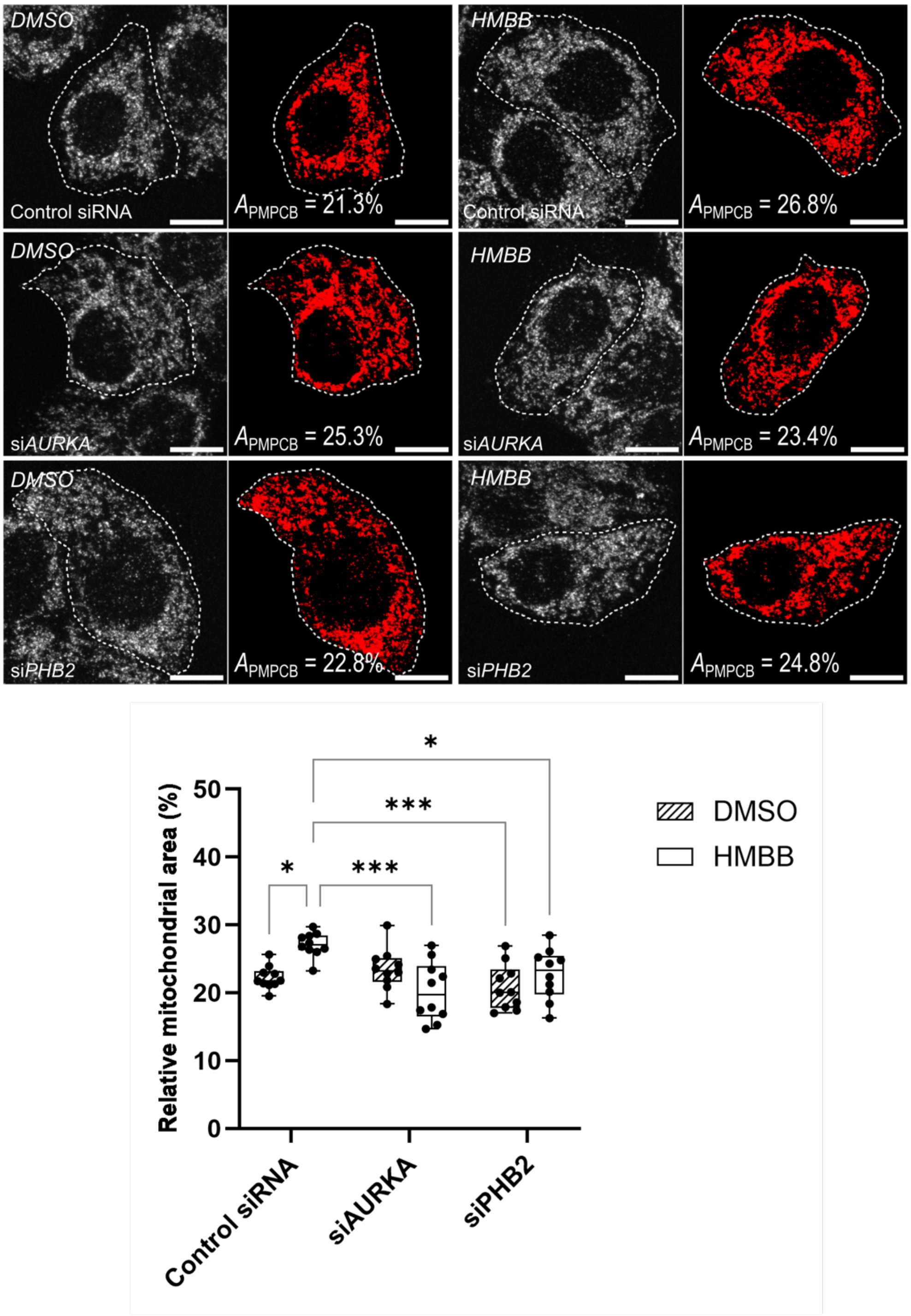


**Supplementary Figure 4**: Mitochondrial mass shown as the amount of PMPCB staining (threshold mask and corresponding quantification) in T47D cells transfected with a control siRNA, or siRNAs targeting *AURKA* or *PHB2* and treated with vehicle (DMSO) or HMBB (50μM). $A_{PMPCB}$: mitochondrial area normalized against total cell area (%). $n$ = 10 cells per condition from one representative experiment (of three). Data are means ± S.D. Scale bar: 10 μm. *$P$<0.05, ***$P$<0.001.

**Supplementary materials and methods**

**CHEMICAL SYNTHESIS**

**General procedure A for the acylation of vanillylamine**

The acyl chloride (3.3 mmol) was added dropwise to a solution of vanillylamine hydrochloride (0.569 g, 3.0 mmol) of sodium hydroxide (0.24 g, 6 mmol) in water (1.2 ml) and THF (1 ml) at 0°C under stirring. After 5 min, the medium was stirred at rt for 2 h, concentrated in vacuo, quenched with 1N HCl and extracted with $Et_2O$. The organic layers were combined, washed with a saturated solution of $Na_2SO_4$, dried ($MgSO_4$) and concentrated under vacuum. The residue was purified by flash chromatography on silica gel using first $Et_2O$ or $Et_2O$/DCM (2:8) and then AcOEt.

**General procedure B for the acylation of vanillylamine**

A mixture of the acid (0.8 mmol), HOBt (108 mg), vanillylamine hydrochloride (151 mg, 0.8 mmol) and triethylamine (82 mg, 0.8 mmol) in 3 ml of anhydrous DMF was cooled down to 0°C under stirring. After 10 minutes, EDCI (173 mg, 0.9 mmol) was added and stirred overnight at room temperature. The medium was the diluted with water, extracted with $Et_2O$ (2X), washed with HCl (1M), a saturated solution of $NaHCO_3$, a saturated solution of $Na_2SO_4$, filtered, concentrated under vacuum. The residue was purified by flash chromatography on silica gel using first $CH_2Cl_2$/pentane 1:1 to 1:0 and then with $CH_2Cl_2$/ $Et_2O$ 8:2 if necessary to afford the expected amide.

***N*-(4-Hydroxy-3-methoxybenzyl)hexanamide (1)**

Following the general procedure A, the amide **1** was obtained in 62 % yield (0.467 g). $^1H$ NMR (500 MHz, $CDCl_3$) δ 6.91 – 6.68 (m, 3H), 5.90 (s, 2H), 4.33 (d, *J* = 5.7 Hz, 2H), 3.86 (s, 3H), 2.27 – 2.10 (m, 2H), 1.70 – 1.55 (m, 2H), 1.30 (d, *J* = 3.6 Hz, 4H), 0.90 – 0.79 (m, 3H). $^{13}C$ NMR (126 MHz, $CDCl_3$) δ 173.16, 146.86, 145.24, 130.43, 120.83, 114.52, 110.82, 56.00, 43.59, 36.88, 31.57, 25.58, 22.49, 14.03.

***N*-(4-Hydroxy-3-methoxybenzyl)octanamide (2)**

Following the general procedure A, the amide **2** was obtained in 34 % yield (0.285 g). $^1H$ NMR (500 MHz, $CDCl_3$) δ 6.93 – 6.69 (m, 3H), 5.79 (s, 2H), 4.35 (d, *J* = 5.7 Hz, 2H), 3.87 (s, 3H), 2.26 – 2.12 (m, 2H), 1.64 (qd, *J* = 7.6, 3.6 Hz, 2H), 1.38 – 1.19 (m, 8H), 0.93 – 0.80 (m, 3H). 13C NMR (126 MHz, CDCl3) δ 173.10, 146.85, 145.25, 130.48, 120.89, 114.51, 110.82, 77.36, 56.03, 43.63, 36.97, 31.80, 29.38, 29.12, 25.91, 22.71, 14.17.

***N*-(4-Hydroxy-3-methoxybenzyl)decanamide (3)**

Following the general procedure B, the amide **3** was obtained in 37 % yield (90 mg). $^1$H NMR (500 MHz, $CDCl_3$) δ 6.91 – 6.72 (m, 3H), 4.36 (d, *J* = 5.6 Hz, 2H), 3.89 (s, 3H), 2.30 – 2.11 (m, 2H), 1.68-1.62 (m, 2H), 1.38 – 1.14 (m, 12H), 0.94 – 0.80 (m, 3H). $^{13}$C NMR (126 MHz, $CDCl_3$) δ 172.98, 146.78, 145.22, 130.53, 120.93, 114.45, 110.78, 77.33, 56.04, 43.64, 37.00, 31.97, 29.56, 29.47, 29.43, 29.37, 25.90, 22.77, 14.21.

***N*-(4-Hydroxy-3-methoxybenzyl)dodecanamide (4)**

Following the general procedure B, the amide **4** was obtained in 52 % yield (140 mg). $^1$H NMR (500 MHz, $CDCl_3$) δ 6.95 – 6.69 (m, 3H), 4.37 (d, *J* = 5.7 Hz, 2H), 3.89 (s, 3H), 2.20 (t, *J* = 7.6 Hz, 2H), 1.66 (p, *J* = 7.4 Hz, 2H), 1.58 (s, 2H), 1.26 (s, 14H), 0.89 (t, *J* = 6.9 Hz, 3H). $^{13}$C NMR (126 MHz, $CDCl_3$) δ 173.01, 146.82, 145.26, 130.58, 120.97, 114.48, 110.81, 56.08, 43.68, 37.04, 32.05, 29.75, 29.64, 29.51, 29.47, 25.94, 22.83, 14.26.

***N*-(4-Hydroxy-3-methoxybenzyl)tetradecanamide (5)**

Following the general procedure B, the amide **5** was obtained in 68 % yield (197 mg). $^1$H NMR (500 MHz, $CDCl_3$) δ 6.89 – 6.69 (m, 3H), 5.63 (s, 1H), 4.37 (d, *J* = 5.6 Hz, 2H), 3.89 (s, 3H), 2.20 (t, *J* = 7.6 Hz, 2H), 1.69 – 1.61 (m, 2H), 1.26 (d, *J* = 3.1 Hz, 20H), 0.89 (t, *J* = 6.8 Hz, 3H). $^{13}$C NMR (126 MHz, $CDCl_3$) δ 173.02, 146.82, 145.26, 130.57, 120.97, 114.48, 110.81, 56.09, 43.68, 37.04, 32.07, 29.82, 29.79, 29.76, 29.65, 29.51, 29.50, 29.48, 25.94, 22.84, 14.27.

***N*-(4-Hydroxy-3-methoxybenzyl)palmitamide (6)**

Following the general procedure B, the amide **6** was obtained in 48 % yield (148 mg). $^1$H NMR (500 MHz, $CDCl_3$) δ 6.92 – 6.72 (m, 3H), 5.61 (d, *J* = 19.9 Hz, 2H), 4.37 (d, *J* = 5.6 Hz, 2H), 3.89 (s, 3H), 2.20 (t, *J* = 7.6 Hz, 2H), 1.66 (p, *J* = 7.5 Hz, 2H), 1.34 – 1.22 (m, 24H). $^{13}$C NMR (126 MHz, $CDCl_3$) δ 173.01, 146.82, 145.25, 130.57, 120.97, 114.48, 110.81, 77.36, 56.09, 43.68, 37.05, 32.08, 29.85, 29.82, 29.80, 29.76, 29.65, 29.51, 29.48, 25.94, 22.84, 14.27.

***N*-(4-Hydroxy-3-methoxybenzyl)stearamide (7)**

Following the general procedure B, the amide **7** was obtained in 81 % yield (270 mg). $^1$H NMR (500 MHz, $CDCl_3$) δ 6.93 – 6.68 (m, 3H), 5.59 (s, 2H), 4.37 (d, *J* = 5.6 Hz, 2H), 3.89 (s, 3H), 2.26 – 2.13 (m, 2H), 1.66 (p, *J* = 7.6 Hz, 2H), 1.26 (d, *J* = 4.0 Hz, 28H), 0.89 (t, *J* = 6.9 Hz, 3H). $^{13}$C NMR (126 MHz, $CDCl_3$) δ 13C NMR (126 MHz, CDCl3) δ 173.02, 146.82, 145.26, 130.58, 120.98, 114.49, 110.81, 56.09, 43.69, 37.05, 32.08, 29.85, 29.83, 29.81, 29.76, 29.65, 29.51, 29.48, 25.94, 22.84, 14.27.

**2-Cyclohexyl-*N*-(4-hydroxy-3-methoxybenzyl)acetamide (8)**

Following the general procedure B, the amide **8** was obtained in 32 % yield (89 mg). $^{1}$H NMR (500 MHz, $CDCl_3$) δ 6.93 – 6.69 (m, 3H), 5.60 (s, 2H), 4.37 (d, *J* = 5.7 Hz, 2H), 3.88 (s, 3H), 2.07 (d, *J* = 7.1 Hz, 2H), 1.84 (d, *J* = 33.1 Hz, 1H), 1.79 – 1.73 (m, 2H), 1.68 (d, *J* = 34.5 Hz, 3H), 1.37 – 1.20 (m, 3H), 1.20 – 1.07 (m, 1H), 1.01 – 0.88 (m, 2H). $^{13}$C NMR (126 MHz, $CDCl_3$) δ 172.27, 146.83, 145.23, 130.62, 120.93, 114.46, 110.75, 77.36, 56.05, 45.14, 43.62, 35.57, 33.34, 26.34, 26.20.

**3-Cyclohexyl-*N*-(4-hydroxy-3-methoxybenzyl)propanamide (9)**

Following the general procedure B, the amide **9** was obtained in 33 % yield (80 mg). $^{1}$H NMR (500 MHz, $CDCl_3$) δ 6.96 – 6.62 (m, 3H), 5.64 (br s, 1H), 5.60 (s, 1H), 4.36 (d, *J* = 5.6 Hz, 2H), 3.89 (s, 3H), 2.18 (t, *J* = 7.6 Hz, 2H), 1.77 – 1.60 (m, 7H), 1.33 – 1.04 (m, 6H). $^{13}$C NMR (126 MHz, $CDCl_3$) δ 173.02, 146.82, 145.26, 130.58, 120.98, 114.49, 110.83, 56.09, 43.68, 37.60, 37.31, 37.20, 33.42, 26.79, 26.75, 26.48, 23.31.

***N*-(4-Hydroxy-3-methoxybenzyl)cyclopentanecarboxamide (10)**

Following the general procedure B, the amide **10** was obtained in 27 % yield (11 mg). $^{1}$H RMN (500 MHz, $CDCl_3$) δ 6.86 (s, 1H), 6.81 (d, *J* = 2.0 Hz, 2H), 6.78 (d, *J* = 2.0 Hz, 1H), 4.37 (d, *J* = 5.6 Hz, 2H), 3.89 (s, 3H), 2.52 (d, *J* = 8.2 Hz, 1H), 1.90 - 1,74 (m, 4H), 1.66 – 1.52 (m, 4H). $^{13}$C RMN (126 MHz, $CDCl_3$) δ 176.13, 146.81, 145.21, 130.70, 120.88, 114.49, 110.75, 56.08, 46.09, 43.68, 30.61, 26.06.

***N*-(4-Hydroxy-3-methoxybenzyl)-3,7-dimethylocta-2,6-dienamide (11)**

Following the general procedure B, the amide **11** was obtained in 52 % yield (252 mg). $^{1}$H NMR (500 MHz, $CDCl_3$) δ 6.94 – 6.70 (m, 3H), 5.64 (d, *J* = 19.7 Hz, 2H), 5.07 (ddddt, *J* = 7.4, 5.9, 4.4, 2.9, 1.4 Hz, 1H), 4.41 – 4.31 (m, 2H), 2.31 – 2.01 (m, 5H), 1.77 – 1.51 (m, 6H). $^{13}$C NMR (126 MHz, $CDCl_3$) δ 167.21, 155.01, 147.06, 145.45, 132.74, 132.66, 130.92, 124.23, 123.57, 121.25, 121.23, 119.45, 118.21, 114.74, 114.71, 111.13, 111.11, 77.62, 56.34, 56.32, 43.67, 41.16, 33.53, 27.06, 26.49, 26.06, 26.03, 25.10, 18.77, 18.08, 18.04.

**(E)-*N*-(4-hydroxy-3-methoxybenzyl)but-2-enamide (12)**

Following the general procedure A, the amide **12** was obtained in 11 % yield (70 mg). $^{1}$H RMN (500 MHz, $CDCl_3$) δ 6.88 (s, 1H), 6.86 (s, 1H), 6.83 (d, *J* = 1.9 Hz, 1H), 6.79 (dd, *J* = 8.0, 1.9 Hz, 1H), 5.80 (dq, *J* = 15.1, 1.7 Hz, 1H), 5.60 (s, 1H), 4.43 (d, *J* = 5.7 Hz, 2H), 3.89 (s, 2H), 1.87 (dd, *J* = 6.9, 1.7 Hz, 2H). $^{13}$C RMN (126 MHz, $CDCl_3$) δ 165.55, 146.58, 145.03, 140.21, 130.16, 124.77, 120.80, 114.24, 110.62, 55.86, 43.44, 17.64.

***N*-(4-Hydroxy-3-methoxybenzyl)butyramide (13)**

Following the general procedure A, the amide **13** was obtained in 68 % yield (457 mg). $^{1}$H NMR (500 MHz, $CDCl_3$) δ 6.87 (d, *J* = 8.0 Hz, 1H), 6.81 (d, *J* = 2.0 Hz, 1H), 6.77 (dd, *J* = 8.0, 2.0 Hz, 1H), 5.67 (s, 1H), 4.36 (d, *J* = 5.6 Hz, 2H), 3.88 (s, 3H), 2.18 (d, *J* = 7.5 Hz, 2H), 1.75 – 1.64 (m, 2H), 0.96 (t, *J* = 7.5 Hz, 3H). $^{13}$C NMR (126 MHz, $CDCl_3$) δ 172.62, 146.58, 145.00, 130.28, 120.68, 114.24 110.56, 55.82, 43.40, 38.63, 19.07, 13.66.

**_N_-(4-Hydroxy-3-methoxybenzyl)butanamide (14)**

Starting with (3,4-dimethoxyphenyl)methanamine and following the general procedure A, the amide **14** was obtained in 16 % yield (113 mg). $^{1}$H NMR (500 MHz, $CDCl_3$) δ 6.83 (s, 3H), 4.39 (d, *J* = 5.7 Hz, 2H), 3.88 (s, 6H), 2.20 (t, J = 7.5 Hz, 2H), 1.70 (sex, J = 7.5 Hz, 2H), 0.97 (t, J = 7.5 Hz, 3H). $^{13}$C NMR (126 MHz, $CDCl_3$) δ 172.60, 149.07, 148.38, 130.95, 119.99, 111.10, 111.06, 55.85, 55.78, 43.31, 38.65, 19.09, 13.68.

**_N_-(2,4-Dimethoxybenzyl)butyramide (15)**

Starting with (2,4-dimethoxyphenyl)methanamine and following the general procedure A, the amide **15** was obtained in 37 % yield (0.264 mg). $^{1}$H NMR (500 MHz, $CDCl_3$) δ 7.19 (d, *J* = 8.1 Hz, 1H), 6.46 (d, *J* = 2.4 Hz, 1H), 6.44 (dd, *J* = 8.1, 2.4 Hz, 1H), 4.37 (d, *J* = 5.7 Hz, 2H), 4.38 (s, 3H), 4.37 (s, 3H), 2.14 (t, *J* = 7.5 Hz, 2H), 1.66 (sex, *J* = 7.5 Hz, 2H), 0.93 (t, *J* = 7.5 Hz, 3H). $^{13}$C NMR (126 MHz, $CDCl_3$) δ 172.35, 160.38, 158.46, 130.51, 118.89, 103.79, 98.49, 55.28, 55.21, 38.77, 38.69, 19.01, 13.62.

**_N_-(4-Aminobenzyl)butyramide (16)**

1-Butyroyl-1H-benzotriazole (0.567 g, 3 mmol) and 4-(aminomethyl)aniline (0.366 g, 3 mmol) were stirred in dichloromethane (10 mL) at room temperature for one night. The solvent was replaced by $Et_2O$. The organic layer was washed with a solution of $K_2HPO_4$ and $NaHCO_3$, dried over $Na_2SO_4$ and concentrated. The oil was purified by column chromatography ($Et_2O$ and AcOEt) and concentrated under vacuum to give the title amide in 37 % yield (0.264 mg) as a pale-yellow powder. $^{1}$H NMR (500 MHz, $CDCl_3$) δ 7.05 (d, *J* = 8.4 Hz, 2H), 6.62 (d, *J* = 8.3 Hz, 2H), 5.81 (s, 1H), 4.29 (d, *J* = 5.5 Hz, 1H), 3.69 (s, 2H), 2.15 (t, J = 7.4 Hz 2H), 1.66 (quint, *J* = 7.4 Hz, 1H), 0.93 (t, *J* = 7.4 Hz, 3H). $^{13}$C NMR (126 MHz, $CDCl_3$) δ 172.57, 145.71, 128.95 (2C), 127.99, 115.00 (2C), 60.22, 43.02, 38.52, 19.01, 13.61.
